# Impact of education and music training on the development of early abstraction abilities

**DOI:** 10.1101/2025.03.28.645996

**Authors:** Morfoisse Théo, Séverine Becuwe, Marie Palu, Cassandra Potier-Watkins, Ghislaine Dehaene-Lambertz, Stanislas Dehaene

## Abstract

Plato’s Republic, Einstein’s Theory of relativity, Vilvadi’s Four Seasons are all remarkable examples of humans’ unique ability to create and manipulate complex abstract structures, whether in language, mathematics or music. Yet the mechanisms by which children develop such abstract thinking, and the role of education and structured experiences such as musical practice in shaping these abilities remain unclear. To explore these questions, we conducted cross-sectional behavioral experiments with 566 children aged 4 to 8, spanning four educational grades, half of whom participated in a violin training program since the age of four. Two experiments examined how children encode, process and compress auditory sequences and visual patterns, while a third examined their sensitivity to geometric regularities. Our results reveal the emergence of symbolic reasoning as early as the start of formal schooling, yet with deeper abstraction as a function of grade. By first grade, children encoded complex auditory sequences within a Language of Thought (LoT) similar to adults. Additionally, when confronted with quadrilaterals, children showed increasing sensitivity to geometric regularities, suggesting a developmental transition from perceptual to symbolic reasoning. However, we did not observe significant impact of musical practice on abstraction abilities across any of the domains tested. We discuss whether and how the impact of education and extracurricular activities such as music could be enhanced.

## Introduction

Whether in the domain of language, mathematics, or music, humans possess a remarkable ability to encode complex information abstractly, by mentally structuring it beyond its linear form (Chomsky, 1965; Dehaene et al., 2015; Hauser et al., 2002). For example, we understand and memorize non-linguistic structures such as auditory or visuospatial sequences by reencoding them internally as nested, tree-like structures using a language-like system (Al Roumi et al., 2021, 2023; Amalric et al., 2017; Chomsky, 1965; Planton et al., 2021; Spelke, 2022; Wang et al., 2019). The *Language of Thought* (LoT) hypothesis has been proposed as one such potential system (Dehaene et al., 2022; Fodor, 1975, 1983; Quilty-Dunn et al., 2023). It stipulates that cognitive representations are constructed from a set of fundamental primitives (e.g., words) that can be recursively recombined through key operations such as concatenation, repetition, and embedding.

Extensive behavioral and neuroimaging studies support this framework. For instance, behavioral studies showed that when participants were asked to memorize geometric sequences drawn on an octagon’s vertices, their error rates were not modulated by sequence length but by its complexity as defined by a LoT-model (Al Roumi et al., 2021; Amalric et al., 2017; Wang et al., 2019) - i.e., the minimum total number of primitive operations required to fully describe them (Chater & Vitányi, 2003; Feldman, 2000, 2003; Mathy & Feldman, 2012). Neuroimaging studies completed the picture, revealing that, rather than memorizing all the individual positions of each sequence items, participants compressed abstractly the information using the sequence’s symbolic structure (Al Roumi et al., 2021; Wang et al., 2019). Similar findings have emerged in auditory tasks, involving memorization and judgment of binary sequences: when the number of items exceeds their working memory capacities (e.g., with sequences of 16 items), adults compressed these sequences into a structured, language-like representation to memorize them (Al Roumi et al., 2023; Planton et al., 2021). Beyond temporal sequences, humans also demonstrate a propensity to encode mathematical and visual objects using abstract, recursive rules (Cheyette & Piantadosi, 2017; Dehaene et al., 2022; Feldman, 2000, 2003; Mathy & Feldman, 2012; Mills et al., 2023, 2024; Piantadosi et al., 2012; Pomiechowska et al., 2024; Sablé-Meyer et al., 2021, 2022). For example, recent works have demonstrated that a visual LoT-model could properly explain human memory for complex geometric shapes such as a square of triangles (Sablé-Meyer et al., 2022), but also the learning of many spatiotemporal patterns, from simple geometrical shapes to intricate mathematical graphs (Mills et al., 2023, 2024).

While the *Language of Thought* framework has shed light on our ability to represent information in a compressed and abstract way, the extent to which it can explain children’s behavior remains uncertain. Some studies suggested that infants exhibit early signs of compositional and logical abilities, which are key components of LoT (Cesana-Arlotti et al., 2018; Feiman et al., 2022; Pomiechowska et al., 2024). Yet, other studies indicated that although these models capture aspects of children’s performance, their explanatory power fall short in comparison to that observed in adults (Mills et al., 2024; Sablé-Meyer et al., 2021). For instance, Sablé-Meyer et al. (2021) found that both an abstract symbolic model and a perceptual convolutional neural network contributed equally to explaining children’s performance on a quadrilateral judgment task. This raises fundamental questions: are these capabilities innate, providing a foundation for acquiring any language, or do they emerge through formal education and experience? While the debate over innate versus learned abilities continues (Cesana-Arlotti et al., 2018; Feiman et al., 2022; Hochmann, 2022; Piantadosi et al., 2018; Pomiechowska et al., 2024), the development trajectory of such abilities and the conditions of education necessary for them to flourish remain underexplored (Alderete et al., 2024; Mills et al., 2024; Sablé-Meyer et al., 2021).

To address these questions, we conducted a series of behavioral experiments with 566 children aged 4 to 8, spanning over four educational grades. This grade range was deliberately selected to capture the developmental shift that occurs before and after the onset of formal schooling – a critical period during which children begin acquiring foundational abilities in areas such as mathematical or reading. Through three experiments designed to assess abstraction abilities, we aim at understanding the impact of formal schooling on these cognitive abilities. These tasks included adaptations of paradigms from prior research on auditory and visual sequences (Planton et al., 2021) and geometric reasoning (Sablé-Meyer et al., 2021), which have successfully demonstrated the relevance of the LoT framework in adults.

Beyond the impact of formal education, we also investigated whether learning another symbolic language – music – enhances abstraction abilities. As we will argue in detail in the next section, since music can be conceptualized as a structured, abstract language, practicing it might foster broader capacities for encoding auditory and visual patterns using a language-like model. We hypothesized that the impact of music education would be maximal on auditory sequences and would diminish as tasks moved farther from the musical domain and towards visual sequences and geometric abstraction. Beyond the three abstraction tasks, we added a fourth experiment measuring inhibitory control, a cognitive function essential for learning (Dehaene, 2018) and which has been reported to be affected by music training (Bolduc et al., 2021; Bugos & DeMarie, 2017; Frischen et al., 2021, 2021; Guo et al., 2018; Holochwost et al., 2017; Janus et al., 2016; Moreno et al., 2011; Shen et al., 2019). This paradigm used is an adaptation of a previously used composite one (Bunge et al., 2002), combining a flanker task (congruent vs. incongruent) (Eriksen & Eriksen, 1974; Fan et al., 2002) with a go/no-go task.

To study this influence of music, the children we recruited from a musical education program, called *Un Violon dans mon école,* implemented in disadvantaged schools of suburban Paris. In half of these schools, children received weekly violin lessons for four years from preschool to second grade. Children in the other half, matched on socio-economic backgrounds, followed the standard curriculum without formal musical instruction. Our sample included 305 children from the violin schools and 261 from the control schools.

### A brief review of the relation between music training and math-related abilities

Music, mathematics, and language all involve the manipulation of abstract, symbolic, and recursive concepts. Some of these concepts may even be shared by several of these fields. For example, the differences between whole notes, half-notes, quarter-notes and eighth notes, which indicate the rhythm in music, echo the mathematical concept of powers of two. Understanding an abstract notion in one of these three domains (e.g., half-notes and quarter notes) could enhance understanding of the same concept in another field (e.g., powers of two). An even stronger hypothesis would be that learning to manipulate abstract notions in one domain (e.g., in music) facilitates the handling of abstract objects in any of these domains (e.g., in mathematics).

At present, however, the evidence suggests that such cross-domain transfer is limited or even inexistent. First, three early longitudinal studies found no significant impact of music training on mathematical abilities (Bilhartz et al., 1999; Costa-Giomi, 2004; Schellenberg, 2004). For instance, Bilhartz et al. (1999), found that 4- and 5-year-olds who were randomly assigned to musical instruction outperformed control groups on visual short-term memory test, but not on formal mathematical assessments. Similarly, Costa-Giomi (2004) followed 117 children over three years and found no improvement in academic achievement, including math, for those taking piano lessons. More recent randomized studies have reinforced these findings, showing little to no effect of music practice on math abilities compared to other artistic activities (Mehr et al., 2013; Rickard et al., 2012). Nevertheless, two other studies (Holmes & Hallam, 2017; Holochwost et al., 2017) provide more nuanced results. In particular, Holochwost et al. (2017) observed that children who participated in an intensive music program (2hours a day) showed improved math and language grades, especially those who stayed in the program for multiple years. These results, however, may have been influenced by the program’s intensity, and the absence of an active control group leaves open the possibility that other intensive activities might yield similar effects. When we shift our focus to general intelligence, the findings remain consistent. Numerous studies comparing music with other activities – such as dance, painting, reading or second-language lessons – report no significant effect of music training on general intelligence (D’Souza & Wiseheart, 2018; Flaugnacco et al., 2015; Janus et al., 2016; Nan et al., 2018). Studies that do suggest weak improvement in intelligence tend to suffer from methodological limitations (Barbaroux et al., 2019; James et al., 2020), have been contested (Schellenberg, 2004; Steele, 2005), or lack control groups (Barbaroux et al., 2019; Osborne et al., 2016).

The only compelling evidence for cross-domain transfer comes from studies explicitly linking mathematics and music, by leveraging the former to teach specific mathematical concepts. For example, a series of studies have demonstrated that the use of music and notions of rhythm can improve children’s understanding of mathematical fractions (An & Tillman, 2015; Azaryahu et al., 2020; Courey et al., 2012; Ribeiro & Santos, 2017). Explicitly connecting two domains during learning therefore seems to be the key to encouraging transfer of knowledge from one to the other (Guilmois et al., 2019; Mayer, 2004; Stockard et al., 2018).

Finally, we note that although the influence of musical practice on mathematics or general intelligence abilities has been examined, general abstraction skills were not specifically assessed with auditory, visual and geometric patterns, as we do here. Yet, these competencies are often viewed as key to the developmental of mathematical understanding and reasoning (Dehaene et al., 2022; Hochmann, 2022; Piantadosi et al., 2016, 2018; Pomiechowska et al., 2024). Whether music learning influences such abilities remains to be demonstrated.

## Materials and Methods

### Participants

292 children, including 115 from violin schools and 177 from control schools, were tested in June 2023. 274 children were also tested in June 2024, with 190 from violin schools and 84 from control schools. Some children tested in 2024 may also have participated in 2023. The children attended 6 nursery and 6 elementary schools, with each level containing three violin schools and three control schools. All schools are in disadvantaged cities in suburban Paris (priority education network with similar social position index). Among the 566 children, 90 were finishing preschool (4-5y old), 165 were finishing kindergarten (5-6y old), 191 were finishing first grade (6-7y old), and 120 were finishing second grade (7-8y old). Ahead of the experiments, all parents received a form to return if they did not wish for their child to participate. Additionally, a consent form was collected from each child before testing began.

### Musical Program

*Un violon dans mon école* is a music education program implemented in disadvantaged French schools. In half of these schools, who volunteered to participate in the program, children received weekly violin lessons from preschool to second grade (4 years). Children in the other half followed the standard curriculum without formal musical instruction, though these schools received budgetary rewards for resources like books. The schools were not randomized at the onset, but they were matched on socio-economic backgrounds. Qualified teachers provided age-appropriate violin lessons, adapting the pedagogy over the four years. Children attended sessions in small groups—45 minutes per week in the first year, increasing to 1 hour 45 minutes thereafter. Lessons took place during school hours, replacing some instruction time.

### Procedure

All children were tested during regular school hours. Four experiments were conducted entirely on a tablet with headphones, lasting 30 to 40 minutes in total. Ten experimenters administered the tests, each overseeing two to three children at a time. Task instructions were pre-recorded in video format to maximize autonomy, with experimenters ensuring proper tablet function and maintaining children’s focus.

## Experiment 1: Auditory abstraction task

Experiment 1 evaluated the development of young children’s ability to detect patterns in auditory sequences and use them to compress information in memory.

### Experimental Paradigm

A previous adult design (Planton et al., 2021) was adapted for children. In this task, two successive eight-note melodies were presented, and children had to judge whether they were identical or whether the second melody deviated from the first by a single deviant note. Each melody lasted two seconds, separated by a one-second silence. Responses were allowed from the onset of the second melody, with an additional two-second window afterward. A one-second pause preceded the next trial. Auditory and visual feedback followed each response to indicate correctness. When present, a deviant note appeared within the last four notes. The melodies followed one of eight patterns, ranging from simple alternations (e.g., ABABABAB) to complex sequences (Figure 1A). The experiment includes 32 trials in total (8 sequences x 2 repetitions with same melodies, and 16 with different melodies).

**Figure 1.**
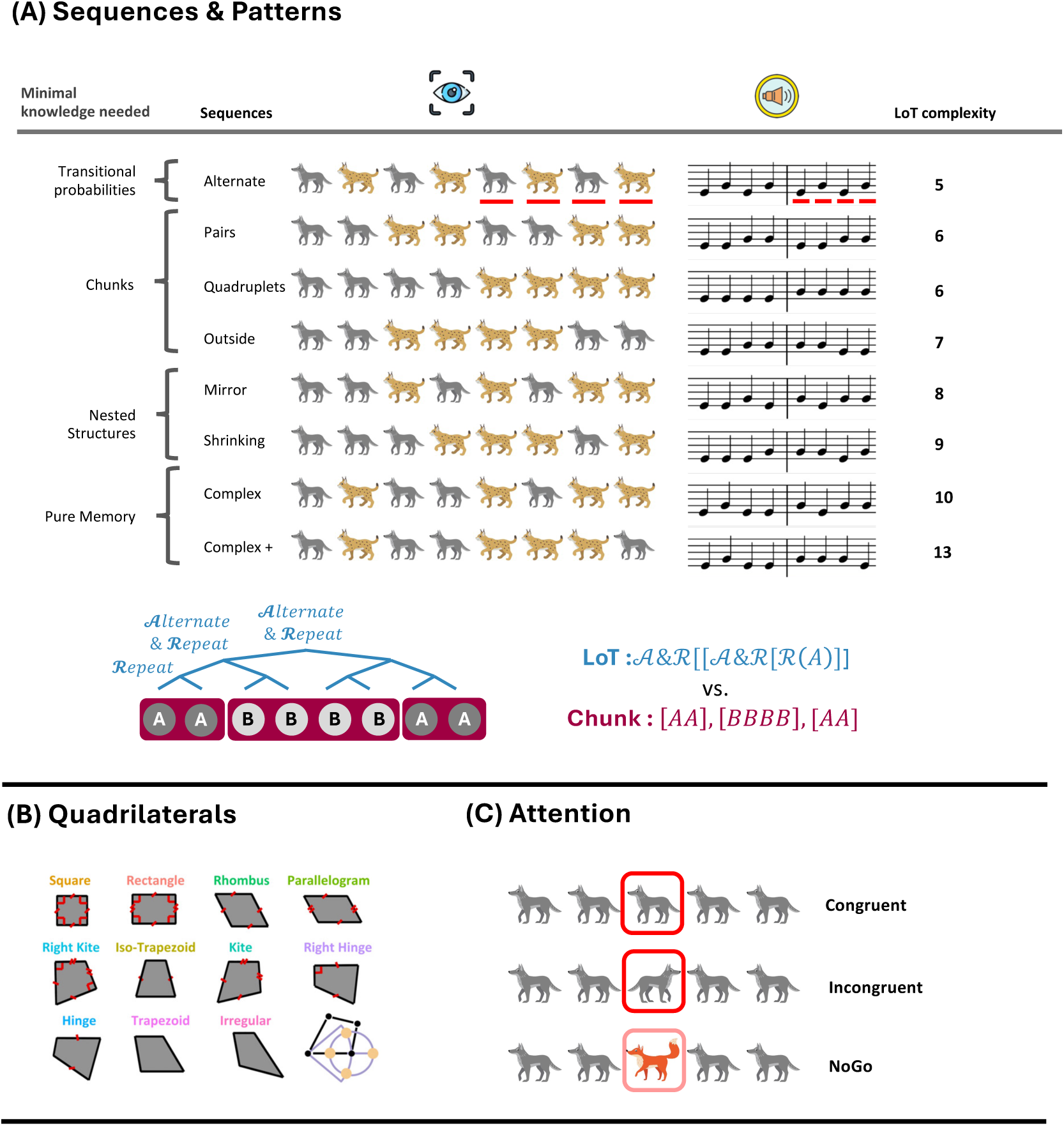
Paradigms. **(A) Abstraction tasks:** children had to judge whether two successive visual patterns (8-animal row) or two auditory sequences (8-note melodies) were identical. Complexity was determined by the *Language of Thought* hypothesis. **(B) Quadrilaterals task:** children had to identify a deviant shape among six quadrilaterals, varying in geometrical regularity (Sablé-Meyer et al., 2021). Deviants were generated by shifting a vertex. **(C) Attention task:** in congruent and incongruent trials, children had to indicate the facing direction of a central animal, while ignoring potential distractors. In no-go trials, children were not supposed to answer.

### Data analyses

We first used one-sample t-tests to determine whether error rates differed from chance levels in each grade. Next, we evaluated various theoretical models from the literature (see below) to explain performance across grades and compared their predictions. For each grade and model, we performed a linear regression between the model’s predicted values and the average performance across participants for each sequence. When necessary, model fit was evaluated using the Akaike information criterion (AIC), which balances goodness of fit against model complexity (i.e., the number of predictors), with the lowest AIC value indicating the best-fitting model. To investigate which model best explained individual performance, we performed linear regressions on each child’s average performance. To compare models at the individual scale, we conducted mixed-model regressions within each grade, comparing the standardized estimates associated with each model with the estimates of all the other models. Participants were included as random effects. Finally, we assessed the potential impact of the music program using a mixed-model binomial regression, with violin training (violin vs. control), grade, and the best-fitting models from the previous analysis as predictors, and participants as a random effect.

### Competing models of sequence encoding

#### Language of Thought (LoT)

In the Language of Thought (LoT) framework, a sequence’s complexity is defined as the minimum number of singular operations needed to fully describe it. These operations, inspired by human language—an embedded and recursive system of thought—include *concatenation* (denoted as “,”), *repetition* (possibly with variations, and denoted as “^n”), and *recursive nesting* (i.e. call of a subprogram). As in human language, a set of primitives which can be recombined recursively using these operations, is required. Only two are necessary here – *stay* (denoted as +0) and *change* (denoted as +1) – since we only have two possible musical notes in each sequence. In this system, a sequence can have multiple possible descriptions. For example, the sequence *AAAA* could be described as a long concatenation of the same element *A*, *A, A, A* (denoted as [+0, +0, +0, +0]), or, in a more compact manner, as the repetition of the same element four times (denoted as [+0]^4). The LoT complexity of a sequence is determined by its shortest possible description in the proposed language (MDL: Minimal Description Length) (Chater & Vitányi, 2003; Feldman, 2000, 2003; Mathy & Feldman, 2012).

#### Chunk complexity

quantifies the degree to which a sequence can be partitioned into subsequences of similar elements, referred to as *chunks*. For example, the sequence *AAABAA* can be divided into 3 chunks – *AAA*, *B*, and *AA*. Mathematically, chunk complexity is expressed by the formula proposed in (Mathy & Feldman, 2012): 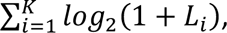 where K represents the total number of chunks and L_i_ denotes the length of the *i*-th chunk. The chunk complexity of the sequence *AAABAA* is then: *log*_2_(4) + *log*_2_(2) + *log*_2_(3).

#### LoT-Chunk complexity

integrates both the Language of Thought (LoT) framework and the concept of chunking. It is defined as the MDL of the shortest possible description in the proposed LoT, but with the additional constraint that the program must respect chunk boundaries. As noted above, a sequence can be described in multiple ways, but some of them do not respect chunk boundaries. For example, the optimal LoT description of *ABBAAB* is [AB] [BA] [AB] (i.e., 3 repetition of the stay-change program; LoT complexity = 5). Yet, this *optimal* description ignores the natural chunks BB and AA, so it is excluded from the definition of LoT-Chunk complexity. The underlying intuition is that humans likely prioritize chunking when processing sequences, before searching for the most compressed representation. LoT-Chunk complexity incorporates this intuition and was found to be the best predictor of adult memory for auditory and visual sequences (Al Roumi et al., 2021; Planton et al., 2021).

#### Lempel-Ziv (LZ) complexity

derives from the Lempel-Ziv algorithm (Lempel & Ziv, 1976), and measures the number of unique substrings encounter during a left-to-right scan of the sequence. For example, in the sequence *AAABAA*, the scan identifies three unique substrings: [A] [AA] [B], resulting in an LZ complexity of 3.

#### Sub-symmetries

quantify the number of symmetric sub-sequences of any length within a sequence (Alexander & Carey, 1968). For example, the sequence *AABBAB* has four symmetric sub-sequences: *AA*, *BB*, *BAB*, and *ABBA*.

#### Change complexity

quantifies the average amount of change across all sub-sequences contained in a sequence, focusing more on the transition from one item to the next, rather than to the items themselves (Aksentijevic & Gibson, 2012).

#### Shannon Entropy

measures the uncertainty associated with a sequence by summing the uncertainties associated of all possible item pairs in the sequence (in our case AA, AB, BA, and BB). Mathematically, the entropy is formulated as:

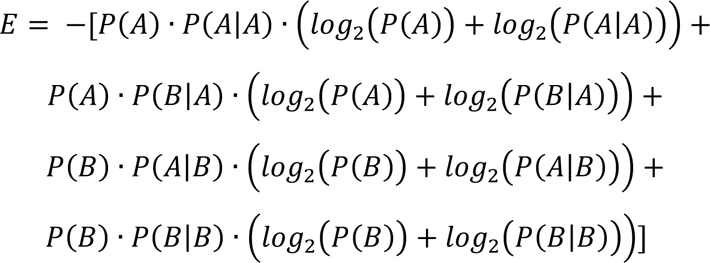

#### Algorithmic complexity (Kolmogorov complexity)

Kolmogorov complexity (KC) is an abstract and non-computable mathematical notion (roughly, the length of the shortest computer program capable of generating a given sequence on a universal Turing machine). Recently, Gauvrit, Delahaye, Zenil and Soler-Toscano (REF) proposed an approximation of KC based on the “coding theorem”, which relates a sequence’s algorithmic complexity to the probability that a universal machine would output it.

Table 1 presents the eight sequences along with the predictor values of their psychological complexity for each model.

**Table 1.** Complexity Scores assigned to each sequence according to the different models.

| Sequences | LoT | LoT-Chunk | Chunk | Change | Entropy | LZ | Algorithmic | Subsymmetries |
| --- | --- | --- | --- | --- | --- | --- | --- | --- |
| ABABABAB | 5 | 5 | 8 | 2.59 | 1 | 4 | 19.84 | 12 |
| AABBAABB | 6 | 6 | 6.34 | 2.51 | 1.96 | 5 | 22.62 | 8 |
| AAAABBBB | 6 | 6 | 4.64 | 1.88 | 1.41 | 5 | 21.9 | 12 |
| AABBBBAA | 7 | 12 | 5.49 | 2.99 | 1.86 | 5 | 22.47 | 10 |
| AABABABB | 8 | 10 | 7.17 | 3.61 | 1.86 | 4 | 22.07 | 8 |
| AAABBBAB | 9 | 9 | 6 | 4.11 | 1.96 | 5 | 22.17 | 8 |
| ABAABABB | 10 | 11 | 7.17 | 3.7 | 1.86 | 5 | 21.58 | 7 |
| ABAABBBA | 13 | 15 | 6.58 | 4.25 | 1.96 | 5 | 21.82 | 7 |

### Results

#### Error rates

Children’s performance gradually improved with grade level (average error percentages of 47%, 43%, 39%, 34% respectively for preschoolers [PS], kindergarteners [KG], first graders [1stG] and second graders [2ndG]). Performance became significantly above chance level from kindergarten onwards (p-values = .11 for PS, < .001 for KG, 1stG and 2ndG; based on one-sample t-tests). Since our task mirrored a similar paradigm in adults with sequences of varying lengths (6, 8, 12 and 16 items) (Planton et al., 2021), we examined the correlation between children and adults. Children’s performance approached that of adults as their grade level increased (r = -.06 and p = .89 for PS; r = .58 and p = .13 for KG; r = .83 and p = .012 for 1stG; r = .78 and p = .022 for 2ndG; see supplementary figure 2.B).

To understand developmental trajectories beyond basic error rates, we evaluated different models proposed from the literature (table 1). Interestingly, a significant linear relationship emerged between LoT complexity and error rates from first grade onwards (r = -.26 and p = .54 for PS; r = .3 and p = .47 for KG; r = .75 and p = .032 for 1stG; r = .87 and p = .005 for 2ndG; supplementary figure 2A). This suggests that as LoT complexity increased, the task of judging similarity became more challenging. While LoT complexity was a strong predictor of error rates, the LoT-Chunk complexity measure provided an even more accurate account of the data. A significant linear relationship was observed between LoT-Chunk complexity and average error rates for first and second graders (r = -.11 and p = .79 for PS; r = .58 and p = .14 for KG; r = .86 and p = .006 for 1stG; r = .94 and p < .001 for 2ndG), which was stronger than the relationship observed for LoT complexity (AIC = 13.98 vs. 14.43 for LoT and LoT-Chunk for PS, 6.12 vs. 3.65 for KG, 15.88 vs. 11.62 for 1stG, and 13.16 vs. 6.5 for 2ndG) (figure 2A). These two LoT-based complexity measures were the best predictors of first and second graders’ performances, outperforming alternative measures such as chunking models, entropy, and algorithmic complexity (figure 2B and E, and supplementary figure 2D). In contrast, none of the complexity models, including probabilistic models, accounted for significant variance in the performance of younger children (i.e. preschoolers and kindergarteners).

**Figure 2.**
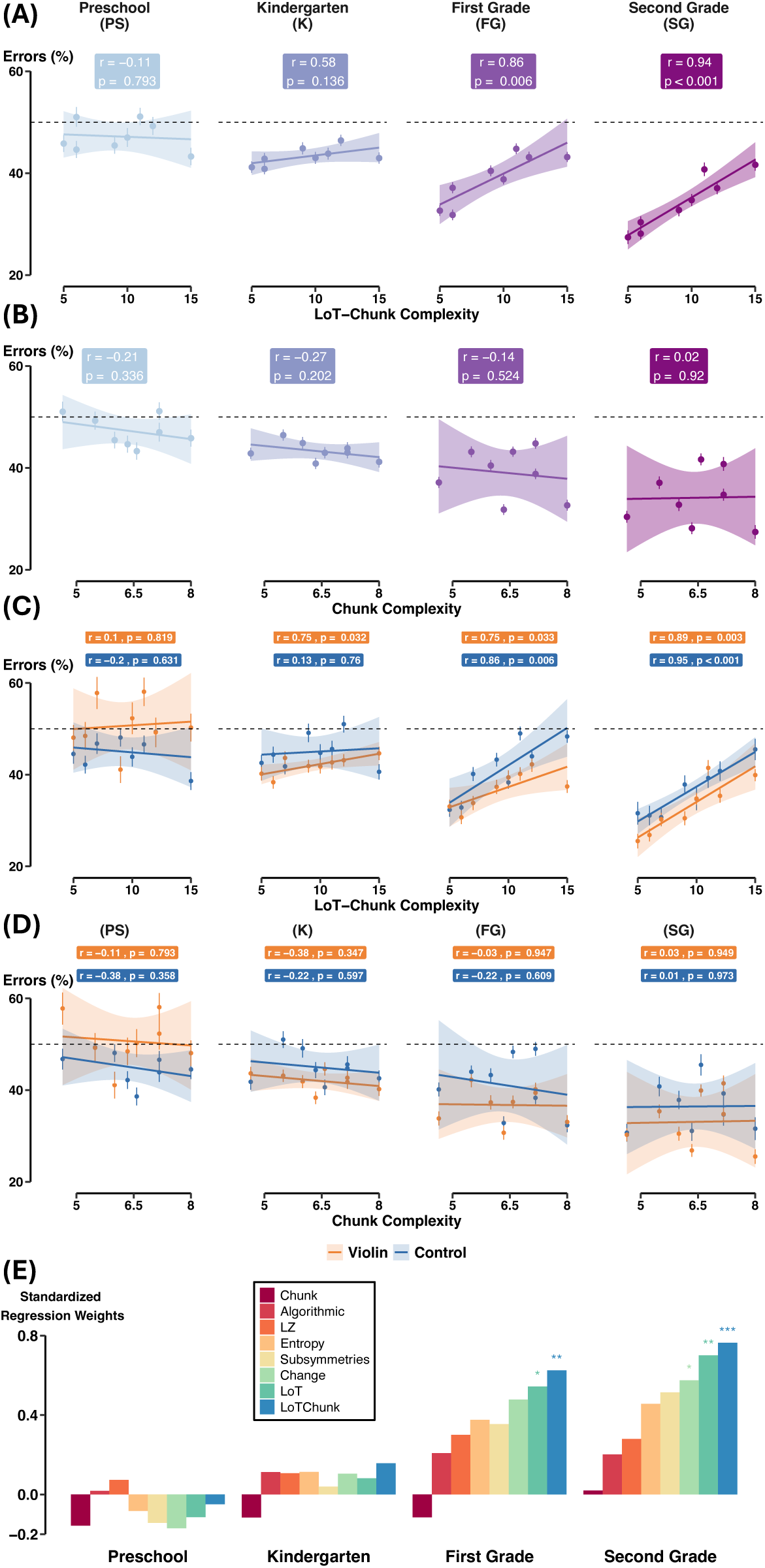
LoT-Chunk Complexity is the best predictor of error rates with auditory sequences. (A-B) Percentage of errors in each sequence, averaged across all subjects within each grade, as a function of LoT-Chunk complexity (A), or Chunk complexity (B). The regression coefficients were referenced in each grade. (C-D) Same as (A-B) but shown separately for violin and control children. (E) Standardized Regression weights from eight linear regressions performed between children’s average performance and model predictions, within each grade.

While the two LoT-related complexity measures were the strongest predictors of average performance in first and second graders, individual differences were observed. Specifically, 30% of first graders and 34% of second graders were best fit by LoT-related models, while 16% and 17%, respectively, were better accounted for the chunking model (only children with at least one R^2^ > 0.2 were included in this best-fit analysis; see supplementary figure 2D for individual estimate). To statistically validate these observations, we conducted mixed-effects linear regressions within each grade, assessing each predictor by comparing its standardized estimates with those of all other predictors. For preschoolers and kindergarteners, no predictor was statically differentiated. For first graders, the chunk predictor’s estimates were significantly lower than all other predictors (p < .001), while LoT and LoT-Chunk estimates were significantly higher (p = .024 for LoT, and p = .014 for LoT-Chunk). A similar pattern was observed for second graders, where LoT and LoT-Chunk again showed significantly higher estimates than other models (p = .0095 for LoT, p = .0016 for LoT-Chunk), while chunk and algorithmic complexity estimates were significantly lower (p < .001 for Chunk, p = .037 for Algorithmic).

#### Response times

RTs increased with grade averaging 2190 ms, 2397 ms, 2620 ms, and 2805 ms, respectively for PS, KG, 1stG, and 2ndG (all trials considered) and 2245 ms, 2451 ms, 2684 ms, and 2828 ms when considering only correct trials (supplementary figure 2C). However, response times were largely uncorrelated with LoT complexity (except surprisingly in KG): r = -.11 and p = .793 for PS; r = .87 and p = .005 for KG and R^2^ = .01 p = .84 for 1stG; r = .41 and p = .32 for 2ndG.

#### Music training

Lastly, we performed a larger mixed-model binomial regression with group (violins vs. controls), grade, LoT-Chunk and Chunk complexities as predictor variables and participants included as a random-effect. Results are presented in table 2, figure 2C and D). Overall, violin students performed slightly better than controls (*p* = .017), with lower average error rates (38.7% vs. 42.1%), and this difference slightly increased with grade level (*p* = .03 for group × grade interaction; Figure 2C). However, when preschoolers were excluded from the regression, the interaction was no longer significant (p = .62), suggesting that the effect was primarily driven—surprisingly—by the poorer performance of preschool violinists (figure 2C). Crucially, the performance of violins and controls varied similarly as a function of sequence complexity (p = .87), and this similarity remained stable across grade levels (p = .45). The same results were observed concerning the interaction between group and chunk complexity (figure 2D).

**Table 2.** Coefficients obtained from a binomial mixed model regression: *Errors* ∼ *Group* ∗ *Grade* ∗ (*LoT* − *Chunk* + *Chunk*) + (1|*Subject*). The predictors Grade, LoT-Chunk and Chunk were standardized prior to the regression.

|  | Estimate | Std. Error | Z-value | P-value |  |
| --- | --- | --- | --- | --- | --- |
| (Intercept) | -0.46 | 0.031 | -14.62 | < 0.001 | *** |
| **Group** | **0.11** | **0.046** | **2.39** | **0.017** | **\*** |
| **Grade** | **-0.22** | **0.031** | **-7.02** | **< 0.001** | **\*\*\*** |
| **LoT-Chunk** | **0.12** | **0.024** | **5.10** | **< 0.001** | **\*\*\*** |
| Chunk | -0.0039 | 0.024 | -0.16 | 0.87 |  |
| **Group x Grade** | **0.099** | **0.046** | **2.17** | **0.030** | **\*** |
| Group x LoT-Chunk | 0.0056 | 0.035 | 0.16 | 0.87 |  |
| Group x Chunk | -0.013 | 0.035 | -0.36 | 0.72 |  |
| **Grade x LoT-Chunk** | **0.067** | **0.024** | **2.77** | **0.0056** | **\*\*** |
| Grade x Chunk | 0.027 | 0.025 | 1.10 | 0.27 |  |
| Group x Grade x LoT-Chunk | 0.026 | 0.035 | 0.75 | 0.45 |  |
| Group x Grade x Chunk | -0.031 | 0.035 | -0.86 | 0.39 |  |

### Discussion

Planton et al. (2021) demonstrated that adults were able to memorize long auditory and visual sequences by using recursive mechanisms to compress them in working memory, as conceptualized by the *Language of Thought* (LoT) framework (Fodor, 1975). Their findings revealed that adult sequence memory performance was strongly modulated by the complexity of the LoT, defined as the minimal number of primitive operations required to describe a sequence. The more complex the sequence, as defined by this language, the more challenging the task of memorization and error detection. When the number of items exceeded their working memory capacities, adults were able to compress these sequences into a structured, language-like representation (e.g., *ABABABAB* as four repetitions of alternating A and B) (Al Roumi et al., 2021; Planton et al., 2021).

The present results replicate and extend those conclusions in children, showing a dual modulation by sequence complexity and participant grade level, with only a minimal influence of music training. Firstly, children’s performance improved significantly with grade and was strongly correlated with the complexity of the sequence. Indeed, once children reached first grade and surpassed chance-level performance (indicating a good understanding of the task), their performance correlated strongly with different models of complexity, and notably with the two complexities related to the Language of Thought hypothesis (i.e., LoT and LoT-Chunk complexities). These two variables were found to be the most predictive of first and second graders’ performance, surpassing alternative models including chunking, entropy, sub-symmetries and more sophisticated models such as algorithmic complexity. This suggests that children at these ages did not memorize sequences simply by partitioning them into chunks, nor by quantifying informative features such as the number or transitions, the probability of transitions, or the number of symmetries or changes. Instead, first and second graders appeared to encode sequences by forming recursive, nested and compressed representations. Moreover, as performance at these ages closely mirrored adult performance on the same task, they seemed to encode auditory sequences very much like adults. These results align with Chomsky’s theories and Fitch’s “dendrophilia” hypothesis, which posits that humans infer hierarchical tree structures from linear sequences even in simple contexts (Chomsky, 1957; Fitch, 2014).

Two further observations are noteworthy. Firstly, while LoT-related complexities were the most predictive models for first and second graders, the LoT-Chunk complexity appeared to be a better estimator than LoT alone, as found in adults (Al Roumi et al., 2023; Planton et al., 2021). This indicates that the optimal description of a sequence should also preserve the chunks in the sequence: humans may first chunk the sequences before compressing them through symbolic operations. Secondly, we explored whether a shift in cognitive strategy occurred over time, with different models being relevant only in younger children and not in older children, and vice-versa. However, none of the models were found to be significant in preschoolers and kindergarteners. Since these younger children struggled to grasp the task, with their performances being around chance level for all sequences, it is difficult to draw any definitive conclusions. Had we employed shorter sequences and extended presentation times, we might have observed a relationship between sequence complexity and performance at these ages. Future studies using implicit, easier, or passive tasks could shed light on the relevance of these models in younger children and offer insights into the developmental trajectory of abstract reasoning abilities prior to school. Previous research already provides evidence of abstraction in preschool-aged children (Piantadosi et al., 2018; Pomiechowska et al., 2024). Notably, Mills et al. (2024) report that 4–6-year-olds can learn to anticipate a wide range of visuospatial sequences, including linear functions and geometrical patterns, with their behavior best predicted by a LoT model—though with a more limited explanatory power compared to adults. Furthermore, given that even 2-year-olds perceive and grasp the syntax of their native language (Brusini et al., 2016), it would be interesting to investigate whether a similar mechanism underlies the abstraction of non-linguistic patterns and (perhaps) contributes to language acquisition.

Beyond the developmental perspective, we explored the influence of musical practice on such abstraction abilities. Like mathematics and language, music involves the manipulation of abstract, symbolic, and recursive structures. This shared characteristic led us to hypothesize that musical training could foster general abstraction abilities, a hypothesis that, until now, has remained largely unexplored.

Our results showed that violinists outperformed control children overall, particularly for children in first and second grade. This suggests that, regardless of sequence complexity, violinists demonstrated a slightly superior ability to memorize and manipulate auditory sequences. This advantage appeared to grow across grades, although this effect was driven solely by the relatively poor performance of the violinists in preschool. However, this benefit of music training could simply reflect a generic enhancement of auditory perception and working memory, which would be helpful for any auditory task, rather than a specific increase in the capacity to compress information using symbolic rules. To differentiate between these two possibilities, we compared the performance of violin and control groups across sequences of varying complexity. The results showed that both groups’ performance evolved similarly with increasing sequence complexity, and that this pattern remained consistent across grades. In other words, both violinists and controls relied on LoT-Chunk-related strategies to the same extent. Thus, while our findings suggest a positive effect of musical practice on auditory processing, we found no evidence that it enhances sequence abstraction abilities. The second experiment, which examines the same question using visual sequences, may help clarify these hypotheses.

## Experiment 2: Visual abstraction task

Experiment 2 evaluated the development of young children’s ability to encode abstract regularities in visual patterns.

### Experimental Paradigm

We used a design similar to experiment 1, but in the visual modality (figure 1A). Two successive fixed animal patterns (8-animal rows) were presented, and children had to decide whether the two patterns were identical. Each pattern was displayed with all animals simultaneously for two seconds, separated by a one-second gray screen. Children could respond either during the second pattern presentation or within an additional 2-second gray screen display. A one-second gray screen then preceded the next trial. Auditory and visual feedback followed each response to indicate correctness. When present, the deviant item was among the last four animals. The arrangement of the animals followed one of eight patterns, ranging from simple alternations (e.g., ABABABAB) to complex sequences (Figure 1A). The experiment includes 32 trials in total (8 patterns x 2 repetitions with same visual rows, and 16 with different visual rows).

### Data Analyses and competing models

The same analyses as in Experiment 1, particularly those exploring the various models, were conducted. These models were the same as in Experiment 1.

### Results

#### Error rates

As in experiment 1, there was a clear developmental progression, with average error percentages decreasing from 50% in PS, 49% in KG, 42% in 1stG, and 31% in 2ndG. Performance was significantly above chance from first grade onwards (p = .1 for PS, p = .78 for KG, and p < .001 for 1^st^ and 2^nd^ graders). However, unlike in the auditory modality, *LoT*-related complexities did not properly account for children’s performance at any grade (figure 3A and supplementary figure 3A). For example, LoT-Chunk complexity correlations were negligible even in the higher grades (all p > .10, figure 3A). Instead, models quantifying pattern features, such as the number or transitions, the probability of transitions, the number of symmetries or of changes, offered better explanations.

**Figure 3.**
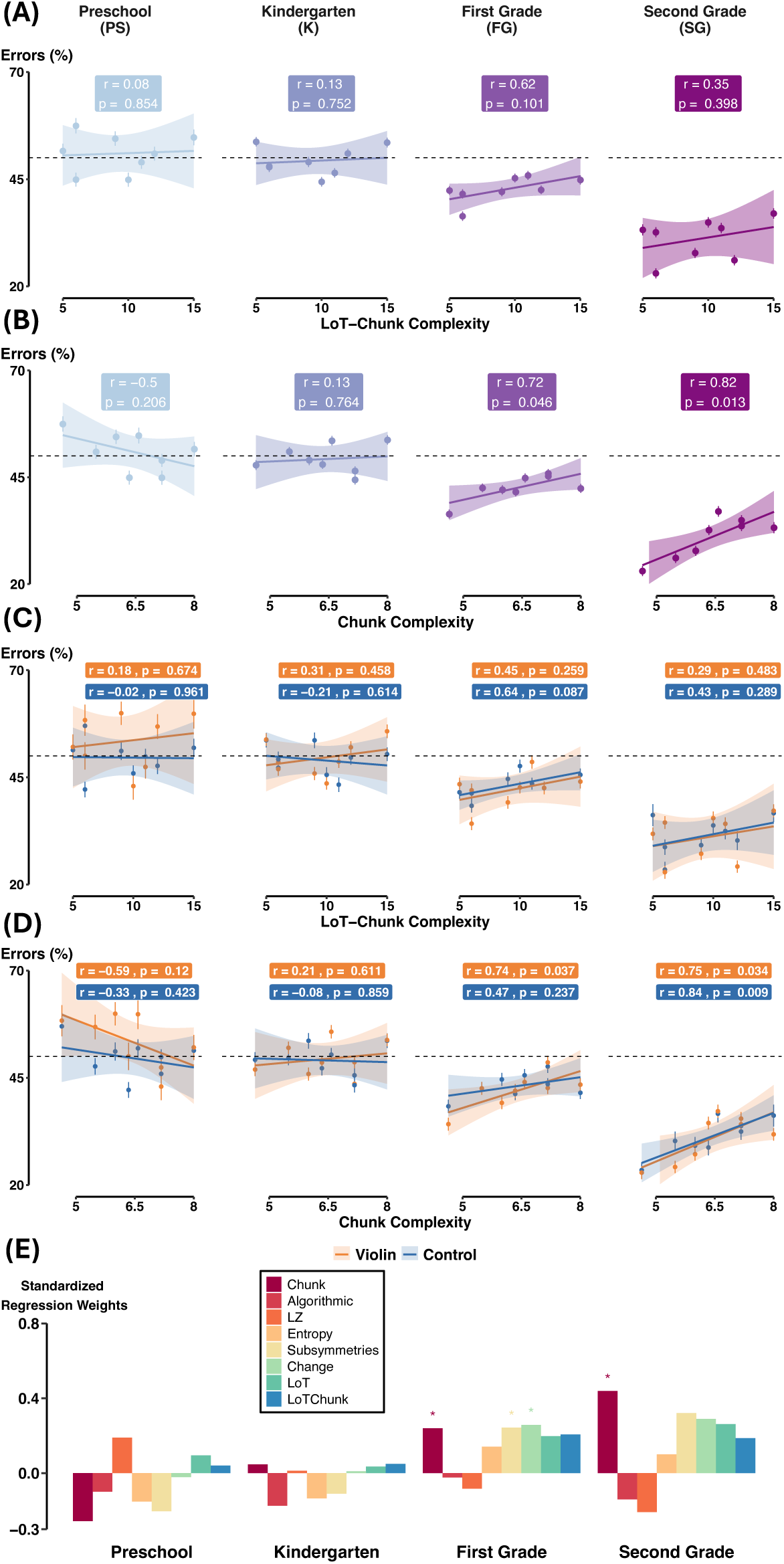
Chunk Complexity, rather than Lot-Chunk, predicts error rates in visual patterns. (A-B) Percentage of errors in each pattern, averaged across all subjects within each grade, as a function of LoT-Chunk complexity (A), or Chunk complexity (B). The regression coefficients were referenced in each grade. (C-D) Same as (A-B) but shown separately for violin and control children. (E) Standardized Regression weights from eight linear regressions performed between children’s average performance and model predictions, within each grade.

The *Chunk model*, which quantifies the extent to which a sequence can be partitioned into subsequences of similar elements, emerged as the most explanatory model for first and second graders: r = -.5 and p = .21 for PS; r = .13 and p = .76 for KG; r = .72 and p = .046 for 1stG; r = .82 and p = .013 for 2ndG (figure 3.B). Nevertheless, the relationship between chunk complexity and performance in the visual modality (e.g., standardized estimate = 0.44 and p = .013 for second graders) was weaker compared to the relationship between LoT-Chunk complexity and performance in the auditory modality (e.g., standardized estimate = 0.76 and p < .001 for second graders). Additionally, none of these models was significant in explaining the performance of younger children (i.e., PS and KG). Figure 3E provides an overview of the standardized regression estimates for all theoretical models examined.

Despite chunk being the most predictive variable on average, individual variability was present, as was the case in the auditory modality. For instance, 18% of first graders were best explained by the chunk model, compared to 16% by the LoT-Chunk model. Among second graders, nearly 30% were best fitted by the same chunk model, while only 9% by the LoT-Chunk model (only children with at least one R^2^ > 0.2 were included in this best-fit analysis; see supplementary figure 3.D for individual estimate). To statistically validate these observations, mixed-effects linear regressions were conducted within each grade group and for each predictor, comparing the standardized estimates associated with each predictor with the estimates of all the other predictors. Results were less conclusive than in the auditory modality, likely due to weaker overall fits across all models in the visual task (as illustrated in supplementary figure 3D). Among preschoolers, only chunk estimates were significantly lower than all others (p = 0.027). For kindergarteners, algorithmic estimates were significantly lower than all other predictors (p = 0.045). In first graders, both algorithmic and LZ estimates were significantly lower than all others (p = 0.009 for Algorithmic and p < 0.001 for LZ). Finally, for second graders, algorithmic, LZ estimates were significantly lower than all others (p < 0.001), while change and chunk estimates were significantly higher than all others (p < 0.001 for Chunk, p = 0.048 for Change).

#### Response times

They increased with grade, averaging 1535 ms for PS, 1462 ms for KG, 1574 ms for 1stG, and 1676 ms for 2ndG across all trials (1525 ms, 1517 ms, 1646 ms, and 1717 ms, respectively, for only correct trials). They were also uncorrelated with LoT complexity (r = -.55 and p = .16 for PS; r = .32 and p = .43 for KG; r = -.53 and p = .18 for 1stG; r = -.41 and p = .31 for 2ndG; supplementary figure 3C).

#### Music training

As in experiment 1, we conducted a larger mixed-model binomial regression using the same predictors – groups (violins vs. controls), grades, Lot-Chunk and Chunk complexities. The results are presented in Table 3. Notably, no significant differences emerged between the violinists and controls (error rates 41.5% for violins vs. 43.5% for controls; figure 3C-D). Furthermore, while violinists demonstrated marginally improved performance with grade in comparison to controls (p = .050), the interaction was no longer significant when the same regression was performed without considering preschoolers (p = .71). As in experiment 1, this interaction was driven by the low performance of violinists in preschool. No interaction between group and other factors reached significance. In other words, the performance of violins and controls evolved similarly as a function of sequence chunk complexity, and this similarity remained stable across grades (figure 3D). The same conclusions applied to the interactions related to LoT-Chunk complexity (figure 3C).

**Table 3.** Coefficients obtained from a binomial mixed model regression: *Errors* ∼ *Group* ∗ *Grade* ∗ (*LoT* − *Chunk* + *Chunk*) + (1|*Subject*). The predictors Grade, LoT-Chunk and Chunk were standardized prior to the regression.

|  | Estimate | Std. Error | Z-value | P-value |  |
| --- | --- | --- | --- | --- | --- |
| (Intercept) | -0.32 | 0.037 | -8.83 | < 0.001 | *** |
| Group | 0.014 | 0.055 | 0.26 | 0.80 |  |
| **Grade** | **-0.35** | **0.037** | **-9.36** | **< 0.001** | **\*\*\*** |
| **LoT-Chunk** | **0.063** | **0.024** | **2.66** | **0.0078** | **\*\*** |
| **Chunk** | **0.097** | **0.024** | **4.03** | **< 0.001** | **\*\*\*** |
| Group x Grade | 0.11 | 0.054 | 1.97 | 0.050 |  |
| Group x LoT-Chunk | -0.0078 | 0.035 | -0.23 | 0.82 |  |
| Group x Chunk | -0.043 | 0.035 | -1.20 | 0.22 |  |
| Grade x LoT-Chunk | 0.025 | 0.024 | 1.03 | 0.30 |  |
| **Grade x Chunk** | **0.091** | **0.025** | **3.67** | **< 0.001** | **\*\*\*** |
| Group x Grade x LoT-Chunk | 0.011 | 0.035 | 0.30 | 0.76 |  |
| Group x Grade x Chunk | -0.032 | 0.035 | -0.91 | 0.36 |  |

### Discussion

Experiment 1 showed that the Language of Thought (LoT) hypothesis effectively predicted performance in the auditory modality. However, in the visual modality, while performance improved with grade, it did not appear to be driven by LoT complexity but rather by the chunk model. How can we explain these differences, given that both experiments used the same pattern hierarchy?

First, it cannot be attributed to poor task performance, as children performed as well on the visual task compared to the auditory one. Second, the difference may stem from the modality itself, as auditory processing is generally superior to vision when it comes to temporal information (Guttman et al., 2005). However, Planton et al., (2021) demonstrated that adult’s performance with sequences of eight items – whether visual or auditory – did align with LoT complexity. They also found a strong correlation between performance across modalities for the same sequences, suggesting a shared cross-model mechanism. Thus, the visual nature of the task alone cannot fully account for these results. However, unlike the auditory task, we presented visual information as a static pattern rather than a sequence unfolding over time. Nevertheless, unpublished data from our lab suggest that even with static visual patterns, adult performance remains influenced by LoT-related measures, though to a lesser extent. We tentatively conclude that, from preschool to second grade, young children seem to rely on simpler models of visual patterns, such as chunk model. This is consistent with a wealth of prior research showing that chunking is widely utilized in memory strategies (Chekaf et al., 2016; Cowan, 2001; Mathy & Feldman, 2012; Miller, 1956).

Beyond the developmental perspective, we also explored the potential influence of musical practice on these abilities. While music was a priori more likely to have an impact on auditory abstraction abilities, we hypothesized that it might also improve visual abstraction, through cross-modal or even amodal mechanisms. We found no differences between groups, suggesting that musical training does not enhance visual working memory. Additionally, there was no significant difference in the impact of complexity, indicating that musical practice has no discernible effect on either abstraction or chunking abilities. This result might confirm that the advantage observed in the auditory modality was probably related to better auditory processing than to a general improvement in sequence abstraction abilities. The third experiment further explores this investigation by examining abstract reasoning in a domain even more distant from music: geometry.

## Experiment 3: Quadrilaterals task

Experiment 3 evaluated the development of young children’s ability to perceive geometric regularities in quadrilateral shapes.

### Experimental Paradigm

This task followed a previous published odd-one-out paradigm (Sablé-Meyer et al., 2021) in which participants had to quickly identify a deviant shape among a set of six. All shapes were quadrilaterals, ranging from highly regular (a square) to fully irregular (an arbitrary quadrilateral devoid of any geometrical properties) (figure 1B). For each reference quadrilateral, four deviant versions were generated by shifting the bottom right vertex by a constant distance, either along the bottom edge or along a circle centered on the bottom-left vertex. In each trial, one of the 11 possible reference quadrilaterals was selected. Five instances of it, varying in scale and orientation (e.g., five rectangles), were presented simultaneously alongside a single deviant (e.g. a rectangle with a displaced bottom-right vertex) on a 3x2 grid. The outlier’s position was randomized, and six levels of shape rotation and shape scale were distributed pseudo-randomly among the six shapes. The participants’ task was to click on the outlier shape as fast and accurately as possible. The experiment includes a total of 44 trials (4 deviants x11 quadrilaterals).

### Competing models

In Sablé-Meyer et al., (2021), the authors contrasted two classes of models of geometric shape perception – a convolutional neural network (CORnet) modeling the ventral visual pathway, and a symbolic model processing nonaccidental geometric properties, such as parallelism and angles. The CNN-complexity of each quadrilateral and its deviants was measured using the L2-norm between their activation vectors from the network’s last layer. In the symbolic model, shapes are represented by vectors of geometric properties (e.g., 1 for right angles, 0 otherwise). The symbolic complexity is then the distance between a quadrilateral’s vector and that of its deviant. More details are provided in the article (Sablé-Meyer et al., 2021).

### Data Analysis

We first used one-sample t-tests to compare error rates to chance levels for each grade. Next, to determine which model best explained children’s data, we performed two-parameters multiple linear regressions within each grade, where the predictors of both models were put in competition. We also performed a mixed-model binomial regression with both models, values from these models and grade as predictor variables, and participants as random effects, to evaluate in greater detail the developmental trajectories of these two models. Finally, we assessed the potential impact of the music program using a mixed-model binomial regression, with violin training (violin vs. control), grade, and the two models as predictors, and participants were included as a random effect.

### Results

Consistent with the previous two experiments, children’s performance improved across grades, with average error percentages decreasing from 70% (PS) to 63% (KG), 61% (1stG), and 52% (2ndG) - all significantly below the chance-level error-rate (83.3%, p < .001) (figure 4A). In line with Sablé-Meyer et al. (2021), the most regular quadrilaterals – *squares* and *rectangles* – stood out in children’s performance. For instance, among kindergarteners, error percentages for squares and rectangles remained below 46%, while exceeding 67% for other more irregular quadrilaterals. This disparity persisted over time, with second graders showing error percentages of 21% and 32% for squares and rectangles compared to over 50% for other quadrilaterals. Despite this dichotomy, average performance was still well predicted by the order of geometrical regularity in all grades (linear regression on 11 points: R^2^ = .29, p = .086 for PS; R^2^ = .45, p = .023 for KG; R^2^ = .54, p = .010 for 1stG; and R^2^ = .61, p = .004 for 2ndG). However, when squares and rectangles were excluded from the analysis (linear regression on only 9 points), the regression was no longer significant (R^2^ = .023, p = .69 for PS; R^2^ = .006, p = .84 for KG; R^2^ = .25, p = .17 for 1stG; and R^2^ = .37, p = .084 for 2ndG). This indicated that the linearity previously observed was mainly driven by the differences in performance between squares and rectangles versus the other quadrilaterals.

**Figure 4.**
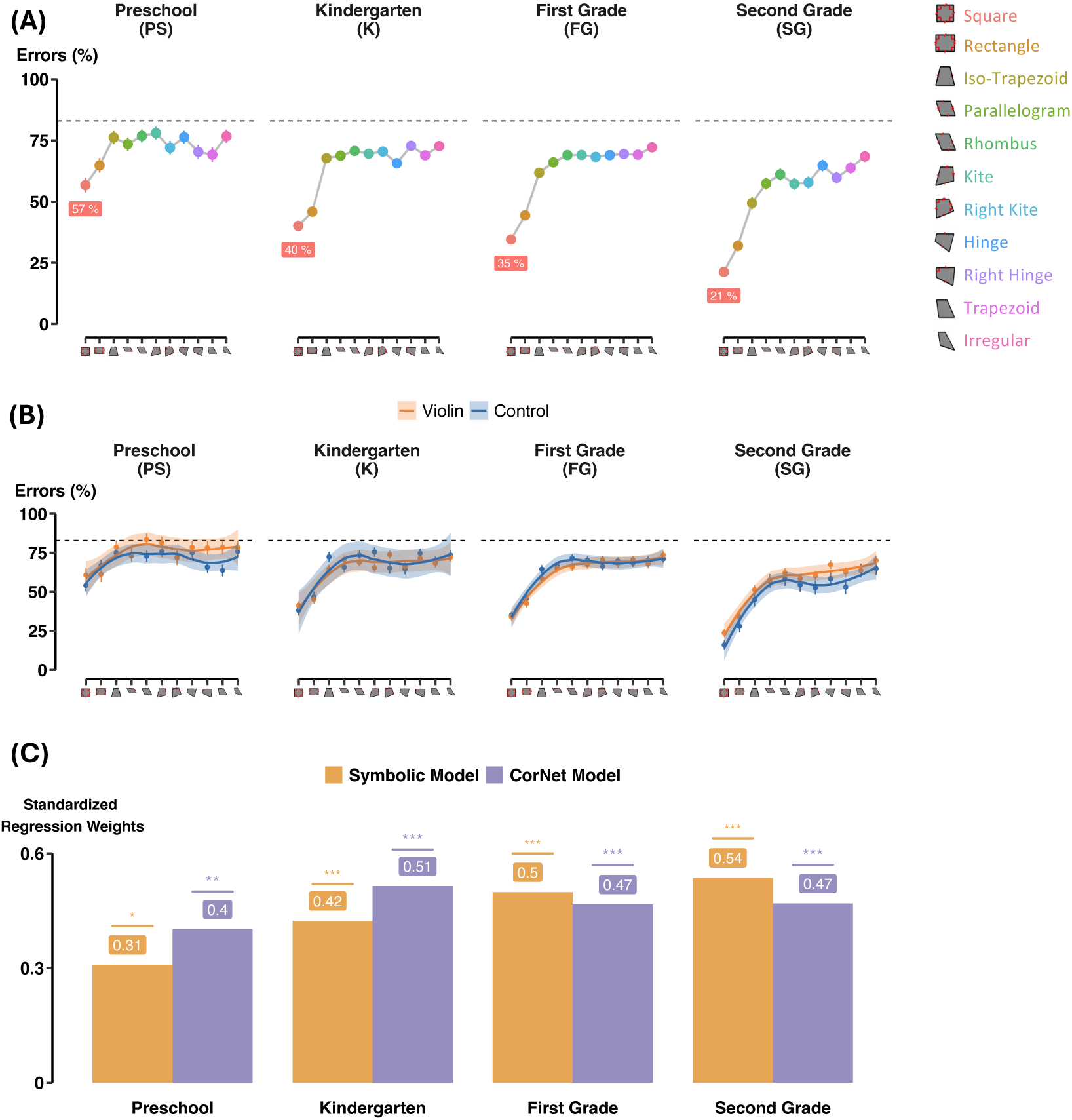
Geometric regularity effect in children. **(A)** Percentage of errors for each quadrilateral, averaged across all subjects in each grade. Shapes are ordered as in Sablé-Meyer et al. (2021). **(B)** Same as (A) but shown separately for violin and control children. **(C)** Standardized regression weights for multiple regressions performed within each grade, across 44 data points (11 shapes x 4 outlier types) using the symbolic and neural network models as predictors. Stars indicate significance level (*•p < 0.1; *p < 0.05; **p < 0.01; ***p < 0.001*).

To better understand developmental trajectories, we examined which of the two models introduced in Sablé-Meyer et al., (2021) – CNNs and symbolic model – best explained children’s performances. Specifically, we performed two-parameters multiple regressions in each grade, where the predictors of both models were put in competition (see figure 4C). Both significantly explained part of the children’s performance (p < .001 for both predictors, for children after preschool). Yet, while correlations with CNNs plateaued over time, those with the symbolic model rose significantly (R^2^ = .31, .42, .50 and .54 for the symbolic model respectively for PS, KG, 1stG, and 2ndG; and R^2^ = .40, .51, .47, .47 for the CNNs). A mixed-model binomial regression with both models, values from these models and grade as predictor variables, and participants as random effects, supported these observations. It revealed that, across successive grades, the symbolic model progressively became a better predictor compared to the CorNet model (p = 0.86 for Symbolic vs. CorNet, and p < 0.001 for Symbolic vs. CorNet x Grade), suggesting that children progressively abandon the shallow visual strategy in favor of an abstract strategy based on geometric rules.

Additionally, we also computed correlations between the performance of children and of the different populations documented in the original article, including Western kindergarteners, first graders and adults, as well as Himba people of northern Namibia - a pastoral community whose language contains no words for geometric shapes and who receive little or no formal education on mathematics - and finally baboons (see Sablé-Meyer et al., 2021, for more details). Two patterns emerged from these analyses (supplementary figure 4). First, across grades, children’s performance became increasingly correlated with those populations, except for baboons with whom correlations remained quite low (R^2^ < 0.45 and decreasing with grade level). Secondly, among KG, 1stG and 2ndG, children’s performance aligned almost perfectly with the children (kindergarteners and first graders) tested in Sablé-Meyer et al. (2021), and to a slightly lesser degree with western adults (e.g. second graders in our study correlated .97, .93 and .75 respectively with kindergarteners, first graders and adults from the original study).

#### Music training

As in experiment 1 and 2, we performed a mixed-model binomial regression with groups (violins vs. controls), grades, and models (Symbolic and CNN) as predictor variables, and participants included as random effects. The results of the mixed-model binomial regression are presented in Table 4. No main effect of group (p = .87, 61.9% errors for violins vs. 63.0% for controls across quadrilaterals) nor interaction of group with other factors was observed (figure 4B).

**Table 4.**
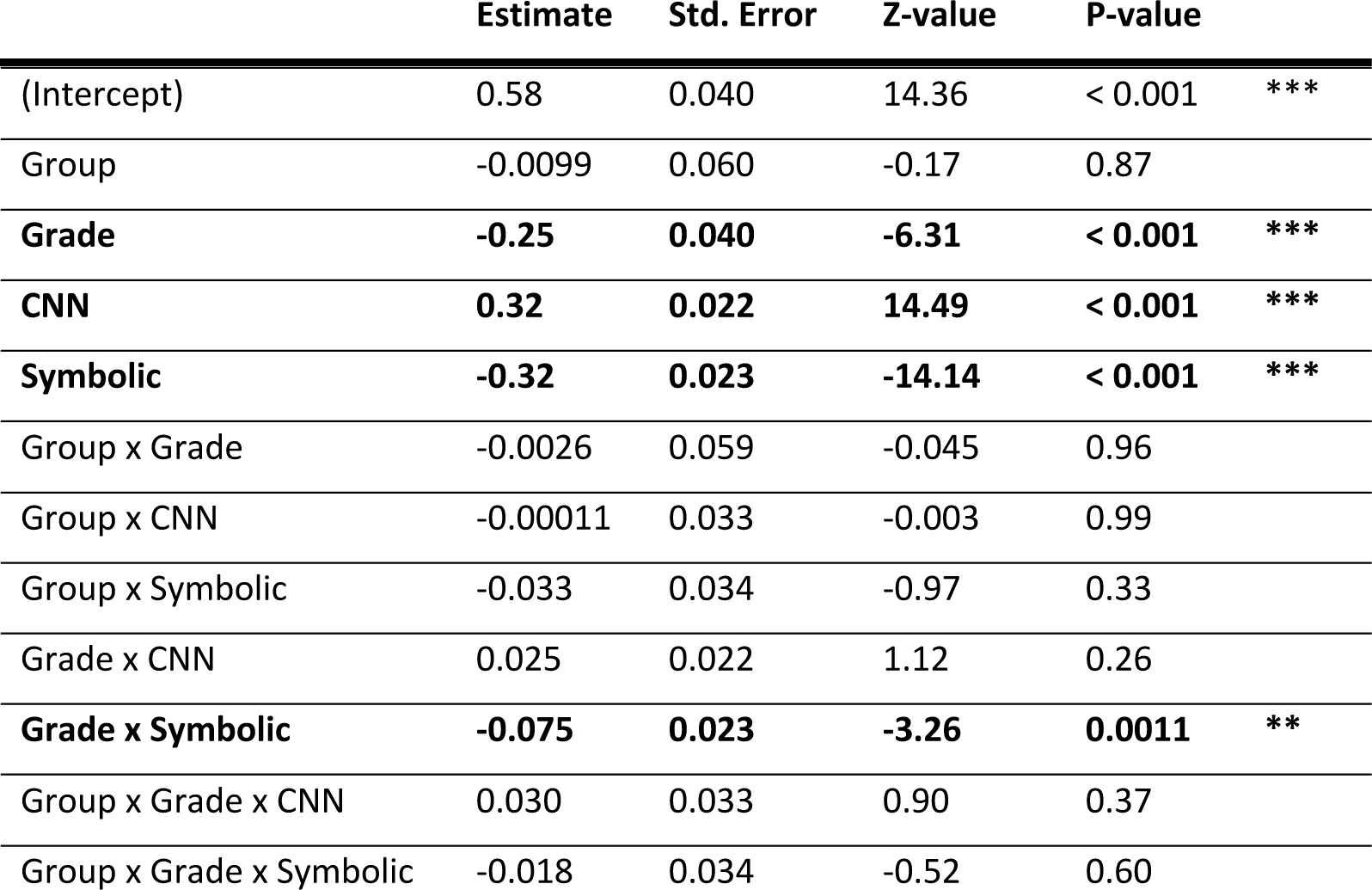
Coefficients obtained from a binomial mixed model regression: *Errors* ∼ *Group* ∗ *Grade* ∗ (*Symbolic* + *CNN*) + (1|*Subject*). The predictors Grade, Symbolic and CNN were standardized prior to the regression.

| Estimate | Std. Error | Z-value | P-value |
| --- | --- | --- | --- |

Table 4. Coefficients obtained from a binomial mixed model regression: $\text{Errors} \sim \text{Group} * \text{Grade} * (\text{Symbolic} + \text{CNN}) + (1|\text{Subject})$ . The predictors Grade, Symbolic and CNN were standardized prior to the regression.
|  |  |  |  |  |  |
| --- | --- | --- | --- | --- | --- |
| (Intercept) | 0.58 | 0.040 | 14.36 | < 0.001 | *** |
| Group | -0.0099 | 0.060 | -0.17 | 0.87 |  |
| **Grade** | **-0.25** | **0.040** | **-6.31** | **< 0.001** | **\*\*\*** |
| **CNN** | **0.32** | **0.022** | **14.49** | **< 0.001** | **\*\*\*** |
| **Symbolic** | **-0.32** | **0.023** | **-14.14** | **< 0.001** | **\*\*\*** |
| Group x Grade | -0.0026 | 0.059 | -0.045 | 0.96 |  |
| Group x CNN | -0.00011 | 0.033 | -0.003 | 0.99 |  |
| Group x Symbolic | -0.033 | 0.034 | -0.97 | 0.33 |  |
| Grade x CNN | 0.025 | 0.022 | 1.12 | 0.26 |  |
| **Grade x Symbolic** | **-0.075** | **0.023** | **-3.26** | **0.0011** | **\*\*** |
| Group x Grade x CNN | 0.030 | 0.033 | 0.90 | 0.37 |  |
| Group x Grade x Symbolic | -0.018 | 0.034 | -0.52 | 0.60 |  |

### Discussion

In Sablé-Meyer et al., (2021), the authors demonstrated that humans, unlike baboons, are sensitive to the regularity of geometric shapes, such as those defined by parallelism, right-angles or symmetries. In the same *odd-one-out* task, where participants had to identify an intruder among six quadrilaterals – one of which has a displaced vertex - adults performed more accurately when the reference shape was a regular quadrilateral (e.g., a square or a rectangle) than when it was a more irregular one, such as a trapezoid. In contrast, baboons showed similar performances regardless of geometric regularity. The latter seemed universal to humans, as it was observed even in a pastoral community with no linguistic terms for geometric shapes and minimal formal education in mathematics, as also in Western preschoolers and first graders.

Our findings replicate and extend these findings, revealing a dual modulation shaped by both geometric regularity and grade, but not by musical practice. Children’s performance improved significantly with grade level and, most crucially, with the geometric regularity of the quadrilaterals. However, although children’s accuracy correlated significantly with the rank of quadrilaterals on the regularity continuum (from square to completely irregular shapes), this relationship was largely driven by the two most regular quadrilaterals – i.e., squares and rectangles. For example, kindergarteners made less than 46% of errors with squares and rectangles, compared to between 67% and 73% with less regular quadrilaterals. The linear relationship between regularity rank and performance vanished when squares and rectangles were excluded from the regression analysis. In previous data from adults, however, this linearity persisted even when excluding those two most regular shapes (Sablé-Meyer et al., 2021). The apparent continuous linearity of the geometric regularity effect might thus reflect a more discrete learning process: children might gradually expand their understanding of regular shapes, progressively incorporating increasingly irregular shapes in their repertoire of known shapes while still considering other shapes as too irregular to be differentiated. Further data from older children will be needed to discern whether this development is continuous or reflects discrete stages. In the latter case, it could suggest that explicit learning, possibly facilitated by learning mathematical vocabulary, is required to fully grasp the concept of geometric regularity.

To explain human and animal performance, Sablé-Meyer et al. (2021) contrasted two classes of models of geometric shape perception: convolutional neural networks (CNNs) that model the ventral visual pathway, and a symbolic model that captures the processing of non-accidental geometric properties such as parallelism and angles. While CNNS effectively captured baboon behavior, only the symbolic model explained human behavior. In our study, although both models explain part of children’s performance, the variance explained by the symbolic system increased significantly with grade, from an R² of .31 to .54, while the explanatory power of neural networks remained essentially constant. This observation suggests that children employed two complementary strategies to perform the intruder task. Initially, they relied mostly on a low-level visual strategy, treating geometric shapes like any other object, a strategy shared by nonhuman primates. Over time, however, children increasingly adopted a more symbolic strategy, interpreting geometric shapes through a set of geometric properties – an approach beyond the reach of current CNNs and nonhuman primates.

While a developmental trajectory was evident, it was not significantly influenced by musical practice. First, the overall performance of children in the violin and control groups was similar across different grades. Second, neither the symbolic nor the CNN model showed stronger correlations with either groups, indicating no significant differences in strategy or performance between musically trained and non-trained children. This result aligns with music training studies and could be explained by the lack of overlap between the domains of geometry and music. Indeed, unlike other mathematical fields, geometry has few, if any, shared notions and concepts with music. The abstract and recursive nature of both domains alone does not appear to be sufficient for musical practice to facilitate the acquisition of geometric abilities. Beyond the concept of abstraction, the final task focuses on examining the more anticipated effects of music on attention abilities.

## Experiment 4: Attention task

Experiment 4 evaluated the development of young children’s attentional selection and inhibition abilities.

### Experimental Paradigm

In each trial, five animals were displayed in a row, and participants had to identify the direction the central animal was facing. If it faced left, they had to press the left button; if it faced right, they had to press the right button. The four surrounding distractor animals (see Figure 1C) influenced the task in different ways. On congruent trials (41.7%), distractors faced the same direction as the central animal, while on incongruent trials (41.7%), distractors faced the opposite direction, requiring participants to ignore them. Finally, on no-go trials (16.7%), distractors were different animals from the central one, signaling participants to withhold their response. To increase task difficulty, no-go trials were deliberately more than twice as rare as the other two trial types.

This task, inspired by a previous study (Bunge et al., 2002), examines two types of attentional control (Posner & Petersen, 1990): *selective attention*, as participants must focus on the central animal (indexed by the difference between incongruent vs. congruent trials), and *executive control* (motor inhibition, indexed by the difference between no-go and other trials).

Each child completed 60 trials: 25 congruent, 25 incongruent and 10 no-go trials. Each trial lasted four seconds during which they could respond, with two seconds of stimulus presentation and two seconds with presentation of a gray background.

### Data analyses

We first used one-sample t-tests to compare error rates to chance levels for each grade. To compare performances and response times between congruent and incongruent trials, we performed mixed-model regressions with condition (congruent vs. incongruent) as predictor variables and participants included as random-effects intercept. We assessed the potential impact of the music program using mixed-model binomial regressions on both performances and reaction times, with violin training (violin vs. control), condition (congruent vs. incongruent) and grade as predictors, and participants included as a random effect. To evaluate performances on no-go trials, we examined d-prime scores, defined as the difference between the z-transformed *hit rate* (responses on congruent and incongruent trials – the go ones) and the z-transformed *false alarm* (responses on no-go trials). Higher d-prime values indicate greater sensitivity. We performed a linear regression on d-prime scores, with groups (violins vs. control) and grade as predictor variables, to assess the impact of music program on executive control abilities.

### Results

#### Error rates

Performance gradually improved with grade level on both congruent and incongruent trials. Specifically, error percentages on congruent trials decreased from 30% to 23%, 12% and 6% and on incongruent trials from 52%, 42%, 28% and 15%, respectively for PS, KG, 1stG and 2ndG. Notably, only the youngest children (PS) failed to perform above chance level on incongruent trials (p < .001 for congruent-trial performance relative to chance level of 50%, and p = .49 for incongruent trials; based on one-sample t-tests within-preschoolers). Across all grades, children performed significantly better on congruent trials than on incongruent ones. This trend was supported by mixed-model binomial regressions performed in each grade, with conditions as predictor variables and participants included as a random-effects intercept (p < .001 for each grade).

#### Response times

Children accelerated with grades for congruent trials (1369 ms, 1338 ms, 1317 ms, and 1273 ms respectively for PS, K, 1stG and 2ndG). In contrast, they becamed slower with grade for incongruent trials (1327 ms, 1395 ms, 1412 ms, and 1396 ms) (supplementary figure 5, A). As a result of this divergence, children from preschool onward responded significantly faster in congruent trials than in incongruent trials, as confirmed by mixed-model linear regressions performed within each group, with conditions (congruent vs. incongruent) as predictor variables and participants included as a random-effects intercept (p = .14 for PS, and < .001 for others).

#### Music training

As in previous experiment, we performed a mixed-model binomial regression with groups (violins vs. controls), grade and condition (congruent vs. incongruent) as predictor variables and participants as random effects. The results are presented in Table 5. No main effect of group (25.0% errors for violins vs. 26.1% for controls) nor interaction of group with other factors was observed (figure 5C), except a slightly significant interaction between group and condition (p = .014). Violinists made marginally fewer errors than controls on incongruent trials compared to congruent trials (averaged across all subjects: 16.8% and 30.7% for congruent and incongruent trials for violins, and 15.7% and 34.6% for controls). Finally, the three-way interaction was not significant (p = .38), indicating no additional differentiation of congruent and incongruent trials in favor of the violin group with grade level.

**Figure 5.**
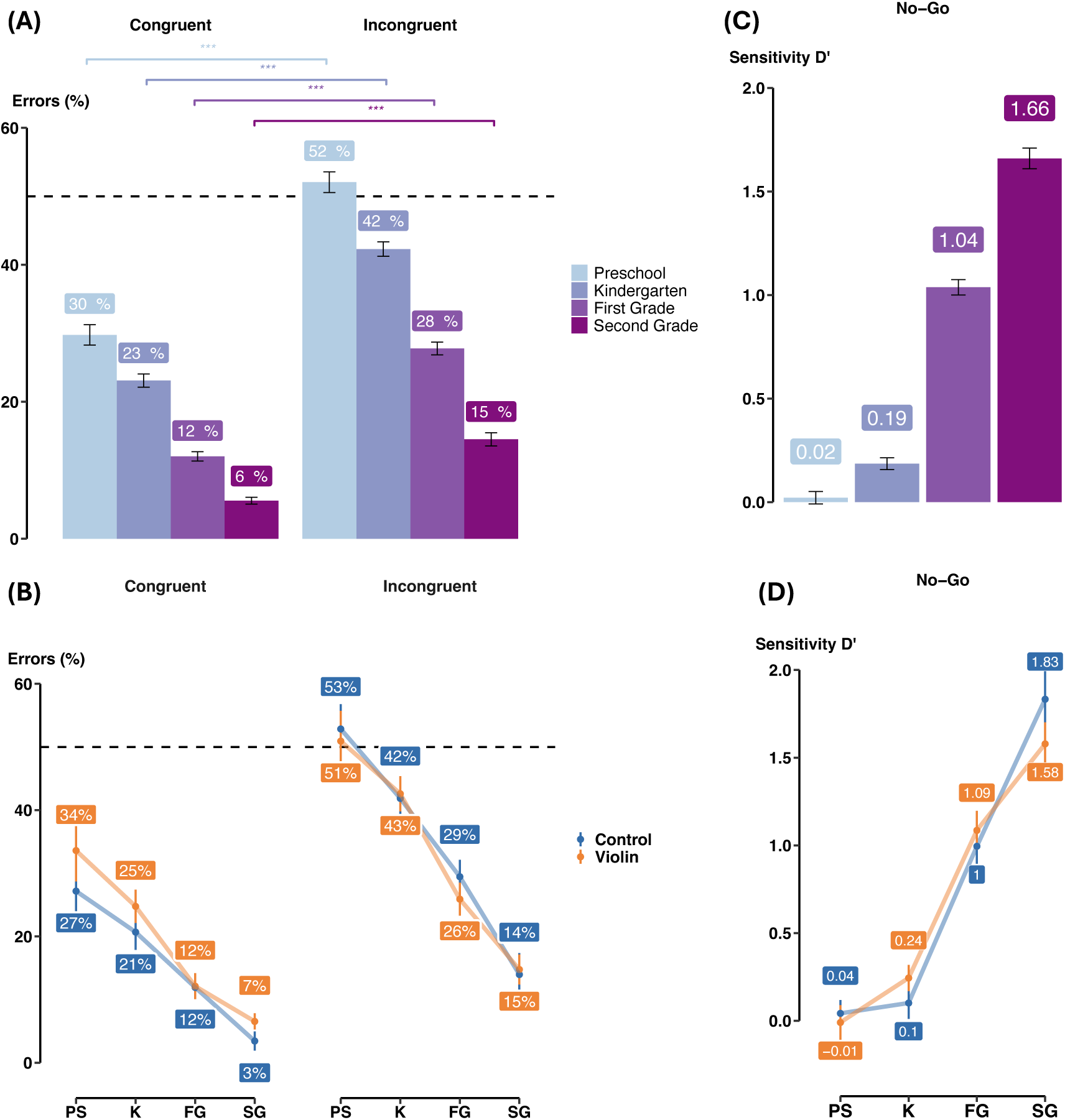
Age, not musical training, enhances the development of attentional abilities. **(A)** Percentage of errors obtained in congruent and incongruent trials, in each grade. Stars indicated significance level from mixed-model binomial regressions performed in each grade: *Errors* ∼ *Condition* + (1|*Subject*). **(B)** D-prime scores, computed as *z*(*hit rate*) − *z*(*false alarm*), within each grade. **(C-D)** Same as (A-B) but shown separately for violin and control children.

**Table 5.**
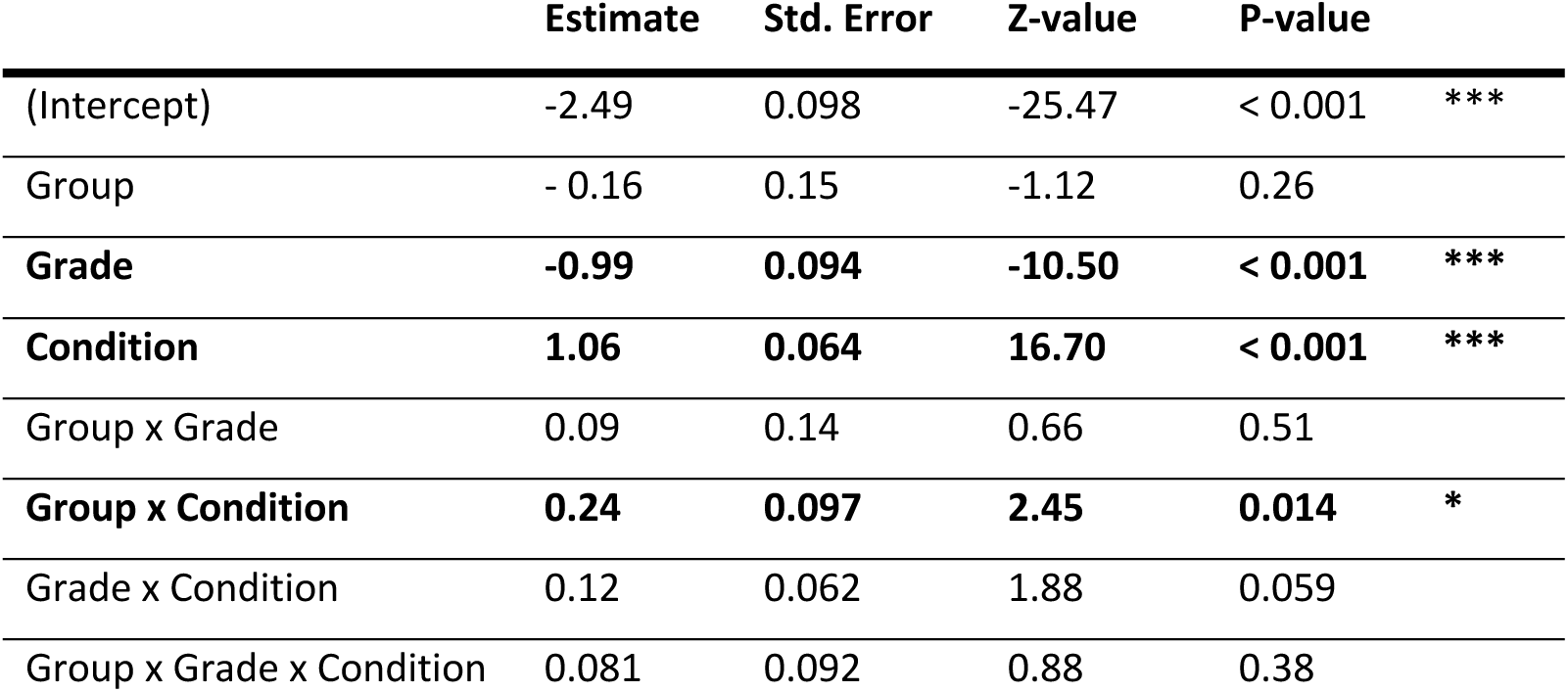
Coefficients obtained from a binomial mixed model regression performed on congruent and incongruent trials: *Errors* ∼ *Group* ∗ *Grade* ∗ *Condition* + (1|*Subject*). The Grade predictor was standardized prior to the regression.

|  | Estimate | Std. Error | Z-value | P-value |  |
| --- | --- | --- | --- | --- | --- |
| (Intercept) | -2.49 | 0.098 | -25.47 | < 0.001 | *** |
| Group | - 0.16 | 0.15 | -1.12 | 0.26 |  |
| **Grade** | **-0.99** | **0.094** | **-10.50** | **< 0.001** | **\*\*\*** |
| **Condition** | **1.06** | **0.064** | **16.70** | **< 0.001** | **\*\*\*** |
| Group x Grade | 0.09 | 0.14 | 0.66 | 0.51 |  |
| **Group x Condition** | **0.24** | **0.097** | **2.45** | **0.014** | **\*** |
| Grade x Condition | 0.12 | 0.062 | 1.88 | 0.059 |  |
| Group x Grade x Condition | 0.081 | 0.092 | 0.88 | 0.38 |  |

The results of a similar mixed-model linear regression performed on response times (only correct responses) are presented in table 6 with no effect related to music training.

**Table 6.** Coefficients obtained from a linear mixed model regression performed on congruent and incongruent trials: *RTs* ∼ *Group* ∗ *Grade* ∗ *Condition* + (1|*Subject*). The Grade predictor was standardized prior to the regression.

|  | Estimate | Std. Error | t-value | P-value |  |
| --- | --- | --- | --- | --- | --- |
| (Intercept) | 1307.14 | 11.31 | 115.55 | < 0.001 | *** |
| Group | -5.80 | 16.96 | -0.34 | 0.73 |  |
| Grade | -10.92 | 11.12 | -0.98 | 0.33 |  |
| **Condition** | **93.18** | **6.99** | **13.32** | **< 0.001** | **\*\*\*** |
| Group x Grade | -3.14 | 16.50 | -0.19 | 0.85 |  |
| Group x Condition | 14.71 | 10.22 | 1.44 | 0.15 |  |
| **Grade x Condition** | **26.30** | **7.11** | **3.70** | **< 0.001** | **\*\*\*** |
| Group x Grade x Condition | 1.97 | 10.55 | 0.19 | 0.85 |  |

#### Executive control (no-go trials)

On average, preschoolers responded on 47% of go trials (i.e., congruent and incongruent ones) and 44% of no-go trials, kindergarteners in 59% of go trials and 51% of no-go trials, first graders in 73% of go trials and 39% of no-go trials, and finally second graders in 82% of go trials, and 29% of no-go trials. To evaluate these performances, we examined d-prime scores, defined as the difference between the z-transformed *hit rate* (responses on congruent and incongruent trials – the go ones) and the z-transformed *false alarm* (responses on no-go trials). As with other measures, children’s sensitivity on no-go trials improved with grade, with d-prime scores (averaged across participants) of 0.02, 0.19, 1.04 and 1.66 respectively for PS, KG, 1stG, and 2ndG (see figure 5.B). Only the d-prime values of the youngest children (PS) were not significantly above zero (p = .72 for PS, .0015 for KG, and < .001 for 1stG and 2ndG; based on one-sample t-tests).

A linear regression performed on d-prime scores, with groups (violins vs. control) and grade as predictor variables, revealed no significant difference between violins and controls (table 7 and figure 5D).

**Table 7.** Coefficients obtained from a regression performed on d-prime scores: *D*′ ∼ *Group* ∗ *Grade*. Only the grade predictor was standardized prior to the regression.

|  | Estimate | Std. Error | t-value | P-value |  |
| --- | --- | --- | --- | --- | --- |
| (Intercept) | 0.77 | 0.053 | 14.40 | < 0.001 | *** |
| Group | 0.014 | 0.078 | 0.18 | 0.86 |  |
| **Grade** | **0.60** | **0.053** | **11.12** | **< 0.001** | **\*\*\*** |
| Group x Grade | 0.021 | 0.079 | 0.27 | 0.78 |  |

### Discussion

We found significant developmental change in both selective and executive attention abilities. The youngest children (preschoolers) made over 50% errors on incongruent trials, while second graders made less than 15%, illustrating a substantial improvement in their ability to inhibit irrelevant peripheral stimuli and focus on the central target. As expected, children across all grades had more difficulty with incongruent trials than with congruent ones, making significantly more errors and responding more slowly. Similarly, children becoming more sensitive, responding more frequently when a response was expected (on go trials, both congruent and incongruent) and withholding more and more responses when none was expected (on no-go trials). The developmental trajectory of these attentional abilities aligns with prior research (Boen et al., 2021; Rueda et al., 2004).

Regarding the comparisons between musically trained and untrained children, we first found a significant, though small, impact on selective attention abilities. Specifically, violin children tended to make marginally fewer errors than controls on incongruent trials compared to congruent trials. However, this interaction may have been largely driven by the surprisingly low performance of violin children on congruent trials (figure 5). Furthermore, while both groups improved with grade level, no one displaying a steeper improvement in performance or response time. The evidence for this slight improvement in visual selectivity therefore remains rather weak, although present, and calls for further research. As regards executive control, there was no difference between the violin and control groups. This absence of results cannot be attributed to a lack of sensitivity, as we easily identified significant differences across grades and suggests that musical practice had no influence on executive control abilities in our sample.

This result may appear surprising, as executive control plays a crucial role in instrumental practice: playing an instrument is a complex motor activity that requires frequent inhibition of automatic, unwanted movements. However, our non-significant results are consistent with the literature. For example, among studies using the Go/No-Go paradigm, only one have demonstrated a clear positive influence of musical learning on executive control (Holochwost et al., 2017; Moreno et al., 2011), and that was an intensive program with two hours of daily training. Other studies have reported mixed (Bolduc et al., 2021; Hallberg et al., 2017; Moreno et al., 2011) or null results (Guo et al., 2018). Similarly, in studies using alternative paradigms, such as the Day/Night Stroop task which requires *verbal* inhibition of automatic responses, only the study without an active control group found positive effects (Bugos & DeMarie, 2017; Frischen et al., 2021; Shen et al., 2019). As for selective visual attention, it has been less frequently examined in this context. One study, using a flanker task, reported positive effects of music training (Holochwost et al., 2017), whereas two other studies employing other selective visual attention tasks, obtained more negative results (James et al., 2020; Janus et al., 2016). Future studies, investigating these different types of attentional selection, including the under-explored area of auditory selective attention, which is perhaps more closely tied to music, could offer new insights into the specific abilities developed through musical training.

## Correlations between experiments

Finally, we examined whether performances in the different tasks were correlated - i.e., whether a child’s performance in one task (e.g., attention) predicted their performance in another task (e.g., quadrilaterals)? To investigate this, we computed the average performance for each participant across the four tasks and calculated the Pearson correlation (defined as 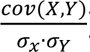) between every pair of tasks. This analysis revealed first that performance on the inhibition task significantly predicted performance on both visual tasks (r = *0.49* and p < 0.001 with abstraction visual; r = *0.42* and p < 0.001 with quadrilaterals). Additionally, performance on the two visual abstraction tasks – patterns and quadrilaterals – was also moderately correlated (r = *0.37*, p < 0.001). By contrast, correlations involving the auditory sequence task were the weakest (r = 0.28 with attention, r = 0.29 with abstraction visual, and r = 0.23 with quadrilaterals; all p < 0.001). Examining development changes across grades, correlation between performance on the attention task and the visual abstract task increased with grade (r= 0.07, 0.25, 0.31, and 0.46, respectively for PS, K, 1stG and 2ndG). However, attention performance did not show a similar trend in predicting performance on the other tasks, related to the language of thought (r = 0.36, 0.34, 0.23 and 0.38, for quadrilaterals; and r = 0.14, 0.15, 0.18 and 0.14, for abstraction audio; across grades). No clear developmental patterns were observed for the correlations between the other task pairs.

While these findings suggest that inhibition abilities positively influence performance on other visual tasks, they do not clarify whether children use similar strategies across tasks. In other words, if a child uses symbolic strategies in one task, does he do the same in another? To address this question, we first computed standardized regression weights for each child in each task, associated to the symbolic models (i.e., LoT-Chunk in auditive and visual abstraction tasks, and symbolic model in quadrilaterals). We then calculated Pearson correlations of these weights between pairs of experiments, separately for each grade. Overall, these correlations were weak or negligible. For instance, there was no correlation between LoT-Chunk estimates from the auditory and visual experiments, indicating that children often used different strategies across these two tasks (as depicted before).

More interestingly, only weak correlations emerged between Lot-Chunk estimates in the auditory task and symbolic model estimates in the quadrilateral task (r = 0.13, p = .20 for second graders). However, these correlations were very low potentially due to a large subset of children not being well-captured by either model. To overcome this limitation, we split children into two groups based on the quadrilateral task: those strongly (i.e., R^2^ > 1) predicted by the CorNet model and those by the symbolic model. For each group, we analyzed whether the standardized regression weights for the LoT-Chunk model differed from those for the Chunk model in the auditory and visual abstraction tasks (see figure 6). The results showed that LoT-Chunk estimates were significantly higher than Chunk estimates only in the auditory task and only for children strongly predicted by the symbolic model in the quadrilateral task (p = 0.024). Indeed, for children better fitted by the CorNet model, no significant differences were observed in this auditory task (p = 0.91), and no significant differences were found for either group in the visual abstraction task (p = 0.69 for CorNet children and p = 0.65 for symbolic children).

**Figure 6.**
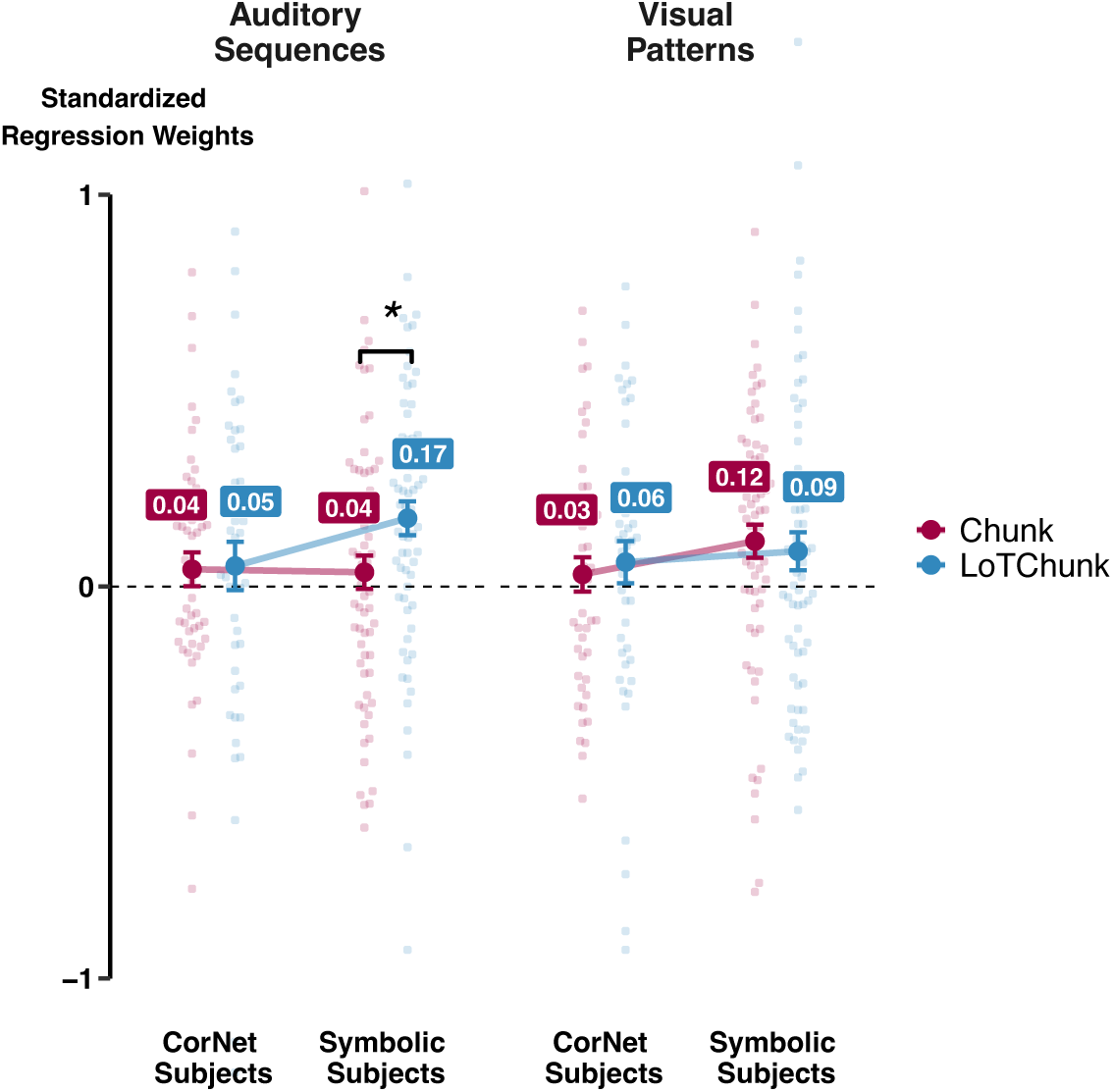
Symbolic strategies common to quadrilateral and auditory sequences tasks. Subjects were separated into three groups: those whose performance in the quadrilateral task was largely predicted by the symbolic model, those by the cornet model, and others. For the first two groups, the regression weights between their performance and predictions of LoT-Chunk and Chunk models in auditory sequences and visual patterns were plotted. Stars indicate whether the regression weights between the two groups of children were significantly different (•P < 0.1; *P < 0.05; **P < 0.01; ***P < 0.001).

### Discussion

By examining between-task correlations, we sought to determine whether children employed consistent strategies across tasks of different nature (e.g., visual patterns vs. auditory sequences). Specifically, we aimed to assess whether children who adopted symbolic strategies in one task (e.g., auditory abstraction), exhibit similar strategies in the two other tasks (e.g., visual abstraction and quadrilaterals). The first striking result was the complete lack of overlap in strategies between the auditory sequence and visual pattern tasks. Children whose performance in the auditory task was well predicted by LoT-related models did not necessarily rely on such models in the visual pattern task. This finding is not surprising given the minimal evidence of LoT-like behaviors observed in the visual pattern task, as noted earlier.

More interestingly, we found a connection between the auditory task and the quadrilateral task. After categorizing children into two groups—those whose performance on the quadrilateral task was best predicted by the CorNet model (*non-symbolic* children) and those best predicted by the symbolic model (*symbolic* children)—we found that, in the auditory task, symbolic children were better predicted by the LoT-Chunk model than by the simpler Chunk model, a pattern not observed in non-symbolic children. In other words, symbolic children were more likely than non-symbolic ones to encode auditory sequences into a structured, language-like representation.

These findings relate to a broader debate on whether there is a single, unified language of thought module or multiple domain-specific modules (Dehaene et al., 2022; Fodor, 1975). Is there one internal mechanism that processes LoT-related operations across all contexts, or are there distinct modules specialized for encoding and compressing structures from different domains, such as visual shapes and auditory sequences? The present results are not conclusive because even if different modules existed for distinct domains, they might nonetheless perform similar recursive operations, allowing a single model to account for behavior across tasks. Thus, the question of a unified versus modular LoT framework remains open for future exploration. Brain imaging during the present tasks may also help, for instance by showing whether task-related activations overlap during music processing, auditory and visual pattern perception, and other mathematical tasks (Al Roumi et al., 2021, 2023; Xie et al., 2022).

## General Discussion

### Development of symbolic reasoning in early childhood

Children between preschool and second grade participated in three experiments to explore the development of their abstraction abilities. By first grade, children began encoding complex auditory sequences using a Language of Thought (LoT), forming hierarchical and structured representations similar to adults. In contrast, visual patterns were processed differently with children relying more on chunking strategies rather than symbolic compression. This suggests either a later emergence of abstract visual processing or a modality-dependent effect. Finally, when confronted with quadrilaterals, children showed increasing sensitivity to geometric regularities, indicating a developmental transition from perceptual to symbolic reasoning.

Across all experiments, particularly experiments 1 and 3, children gradually moved from basic perceptual grouping to structured, rule-based thinking, underscoring the progressive development of abstraction cognition. These findings contribute to a growing body of research highlighting children’s early ability to encode both linguistic and non-linguistic information in abstract ways (Alderete et al., 2024; Cesana-Arlotti et al., 2018; Ciccione et al., s. d., 2023; Feiman et al., 2022; Mills et al., 2024; Pomiechowska et al., 2024; Sablé-Meyer et al., 2021).

### Impact of musical practice

We also wanted to investigate the impact of musical practice, as proposed by the French educational program *A Violin in my school*, on the development of attention and abstraction abilities. These experiments were initially designed to evaluate abilities which are theoretically further and further away from those that are explicitly relevant for music training: starting with attentional abilities, then auditory and visual abstraction abilities, and finally geometrical reasoning.

Our results suggest that musical practice, as implemented in this program, had no significant influence on abstraction abilities, regardless of its precise nature (auditory sequences, visual patterns or geometric shapes). In other words, the abstract, symbolic and recursive nature of music does not seem to be sufficient to facilitate the acquisition of general abstraction abilities. These null results provide explanations for previous studies demonstrating no effect of musical practice on mathematical ability. Indeed, abstraction competencies are often seen as key to the development of mathematical understanding and reasoning, and at the origin of our mathematical abilities. The absence of any effect on such abilities could then explain why, in the absence of a direct link made during learning (An & Tillman, 2015; Azaryahu et al., 2020; Courey et al., 2012; Ribeiro & Santos, 2017), musical practice does not facilitate the acquisition of mathematics (Bilhartz et al., 1999; Costa-Giomi, 2004; Mehr et al., 2013; Rickard et al., 2012; Schellenberg, 2004). However, one might argue that musical education offered to children aged 4 to 8 in this program does not involve enough complex, symbolic and abstract concepts. The program is primarily an introduction to music, focusing on practical musical skills like rhythm and only superficially addressing musical notation. These symbolic aspects typically emerge later in the process of learning to play a musical instrument, with more advanced training in music reading, music theory and solfeggio. An intervention targeting older children might yield different outcomes. Unfortunately, few intervention studies with older children or adults have explored the connection between music and mathematics. Those that exist have not produced clear or generalizable conclusions (Holochwost et al., 2017; Rickard et al., 2012), as argued in the introduction. Furthermore, although numerous cross-sectional studies and general folklore associate mathematical and musical abilities (Cheek & Smith, 1998; Rauscher et al., 1993; Schmithorst & Holland, 2004; W D Ross, 1937), a causal link may be non-existent, due to a third variable, or even run the other way. For example, a correlational study concludes that the correlation between musical practice and cognitive abilities such as math “appears to arise because high-functioning individuals with high levels of musical ability are also more likely to take music lessons” (Schellenberg & Lima, 2023; Swaminathan et al., 2017).

While our results challenge, once again, the existence of a far transfer between music and non-musical domains, they do not rule out the possibility of transfer to closer areas or more transversal skills such as auditory working memory or attentional functions. Indeed, findings from experiment 1 revealed a modest advantage of musical practice on auditory working memory. In contrast, however, findings from experiment 4 suggested little or not effect on attention. There was only a slight enhancement in visual selectivity (though the evidence remains weak and calls for further investigation) and no measurable effect on executive control. These results fall within a still unclear line of research (Schellenberg & Lima, 2023), with some studies showing a clear benefit of musical learning on the development of executive functions (Bolduc et al., 2021; Holochwost et al., 2017; Moreno et al., 2011; Shen et al., 2019), and others demonstrating minimal (Frischen et al., 2021) or no effects (Bugos & DeMarie, 2017; Frischen et al., 2019; Guo et al., 2018; Hallberg et al., 2017; James et al., 2020; Janus et al., 2016; Linnavalli et al., 2018). The fact that these studies vary widely methodologically (e.g., in samples sizes, group randomization or the specific executive functions tested) makes the generalizations of the results highly challenging and uncertain (as depicted in supplementary figure 1B). However, recent reviews have concluded that studies using randomized designs with active control groups did not find evidence that musical training improves executive functions (Schellenberg & Lima, 2023).

In fact, this observation is not limited to executive functions. As described in recent meta-analyses and literature reviews (Bigand & Tillmann, 2021, 2022; Sala & Gobet, 2017, 2020; Schellenberg & Lima, 2023), studies without active control groups are more likely to report significant effects. Although the present program lacks a clearly defined active control group, it is worth noting that violin lessons in this program were conducted during school hours, meaning that control children had more time dedicated to traditional schooling. In both cases, our experiments were not directly testing school-taught knowledge, but rather strategies to resolve novel problems, where the influence of schooling vs brain maturation remains unclear. Future studies should include more well-defined active control to better determine whether musical training influences inhibition and abstraction abilities.

Despite this limitation, our study stands out positively in several ways. First, it includes a larger dataset than most similar studies, with 566 participants compared to the average of 130 (ranging from 71 to 256) in studies on musical practice and mathematics for example (see supplementary figure 1A). Second, the music program was frequent and intense, with over 150 hours of instruction spanning four school years. Finally, our study allows us to measure the cumulative impact of music training by evaluating children after 1, 2, 3 or 4 years of music lessons. These factors give us confidence in concluding that, under the conditions in which it was provided here, musical practice has no measurable effect on general abstraction and only small impact on attention abilities.

To move beyond this disappointing situation, two key factors should be considered. First, it might be more effective to schedule music lessons at a different time, ensuring they do not reduce the standard school curriculum. Second, the literature suggests that for music to positively impact mathematical abstraction abilities, its concepts should be explicitly integrated into instruction - for instance, highlighting the connections between musical notes, fractions, and powers of two. Once again, the present study reinforces that without guidance and explicit instruction, discovery learning has little impact on young children’s understanding of abstract mathematical rules and patterns (Guilmois et al., 2019; Mayer, 2004; Stockard et al., 2018).

## Acknowledgments

This research was funded by the Vareille Foundation. We sincerely thank Fosca Al Roumi, Samuel Planton and Mathias Sablé-Meyer for their valuable input in adapting their paradigms and analyses. We also thank Claire Njoo Deplante for her useful suggestions on the manuscript. We are also grateful to the children who participated.

## Supplementary Figures

**Supplementary Figure 1.**
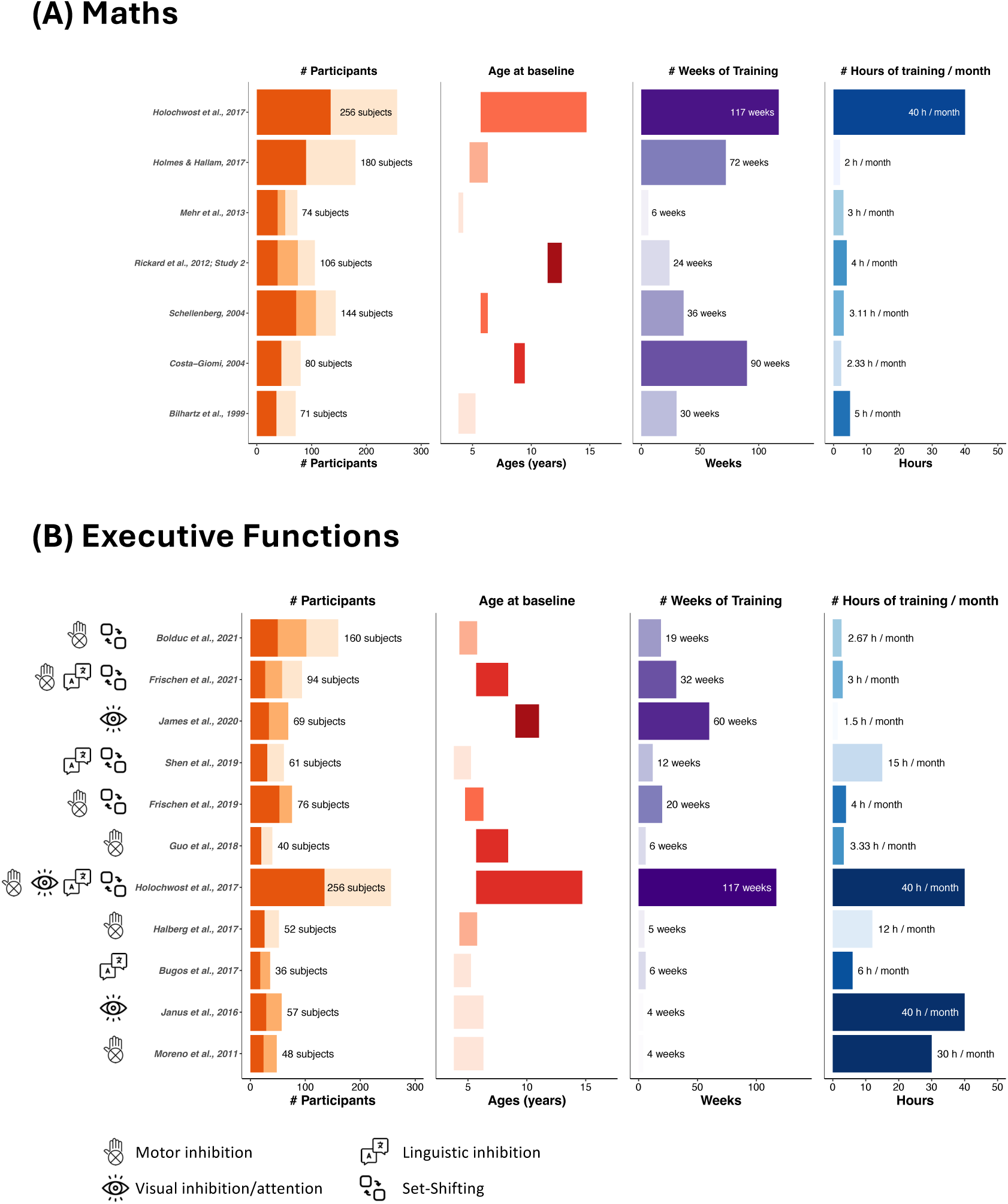
Review of previous articles assessing the impact of musical interventions on the development of children’s mathematical (A) and attentional (B) capacities.

**Supplementary Figure 2.**
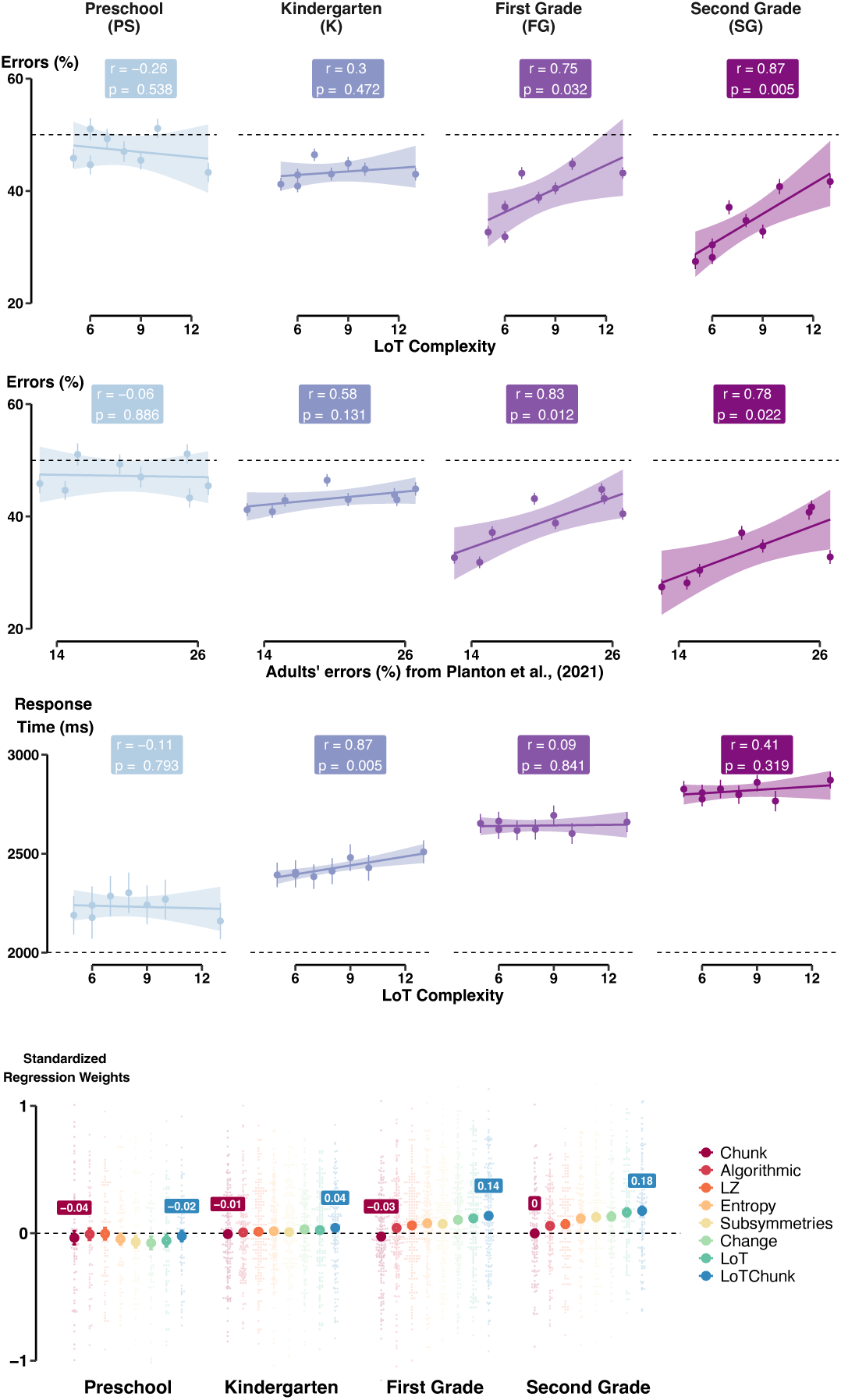
Auditory sequences. (A-B) Percentage of errors in each sequence, averaged across all subjects within each grade, as a function of LoT complexity (A), or adult’s errors as referenced in Planton et al., 2021 (B). The regression coefficients were referenced in each grade. (C) Same as (A) but for response time. (D) Standardized regression weights obtained for each child from eight linear regressions between his/her performance and the model predictions, separately in each grade.

**Supplementary Figure 3.**
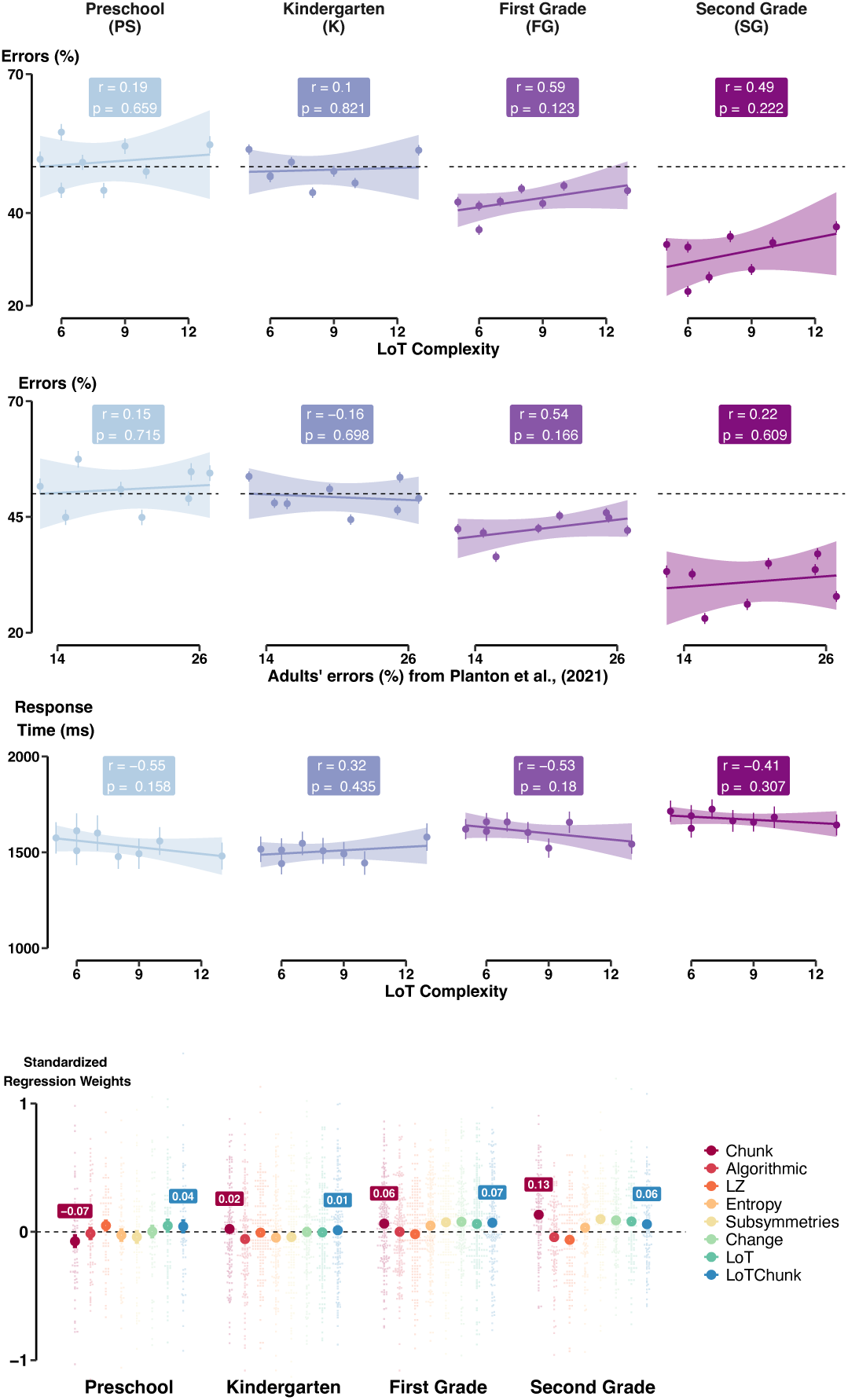
Visual patterns. (A-B) Percentage of errors in each pattern, averaged across all subjects within each grade, as a function of LoT complexity (A), or adult’s errors as referenced in Planton et al., 2021 (B). The regression coefficients were referenced in each grade. (C) Same as (A) but for response time. (D) Standardized regression weights obtained for each child from eight linear regressions between his/her performance and the model predictions, separately in each grade.

**Supplementary Figure 4.**
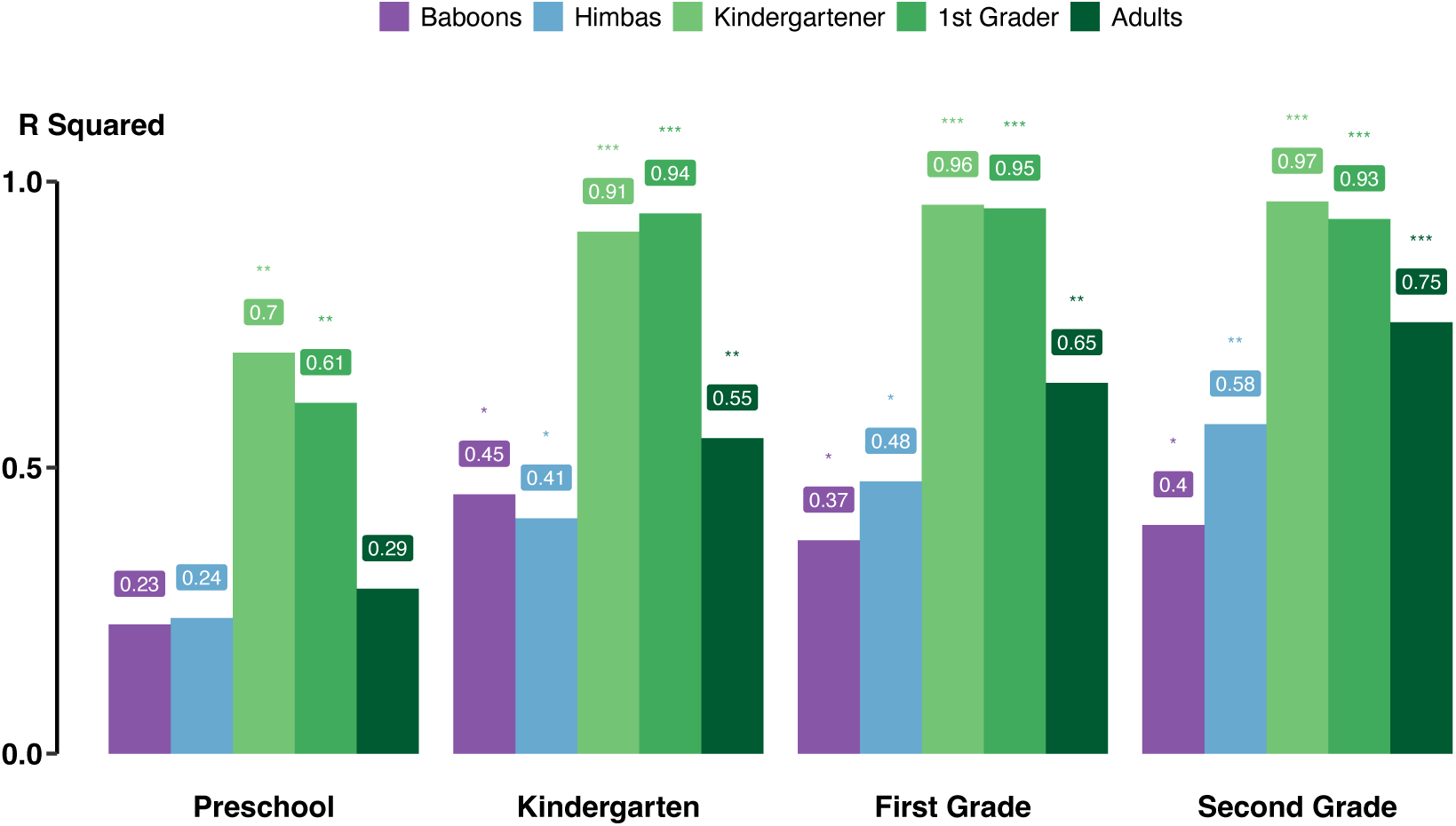
Quadrilaterals. Correlations between the children’s performances and those of the different populations documented in the original article (Sablé-Meyer et al., 2021), including Western kindergartener, first graders, adults, Himba people and baboons.

**Supplementary figure 5.**
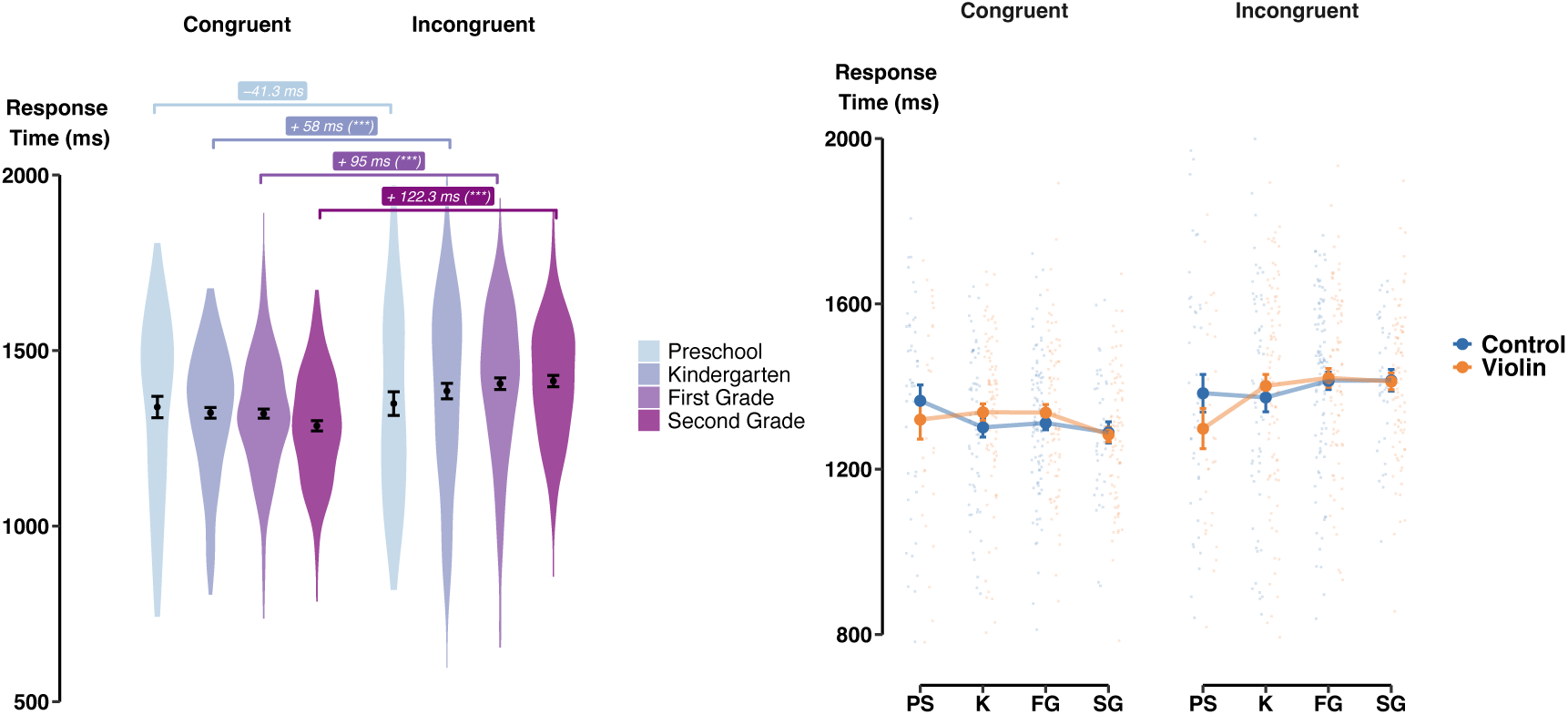
Attention. **(A)** Response Times in congruent and incongruent trials, in each grade. Stars indicated significance level from mixed-model binomial regressions performed in each grade: *RTs* ∼ *Condition* + (1|*Subject*). **(B)** Same as (A) but shown separately for violin and control groups.

**Supplementary Figure 6.**
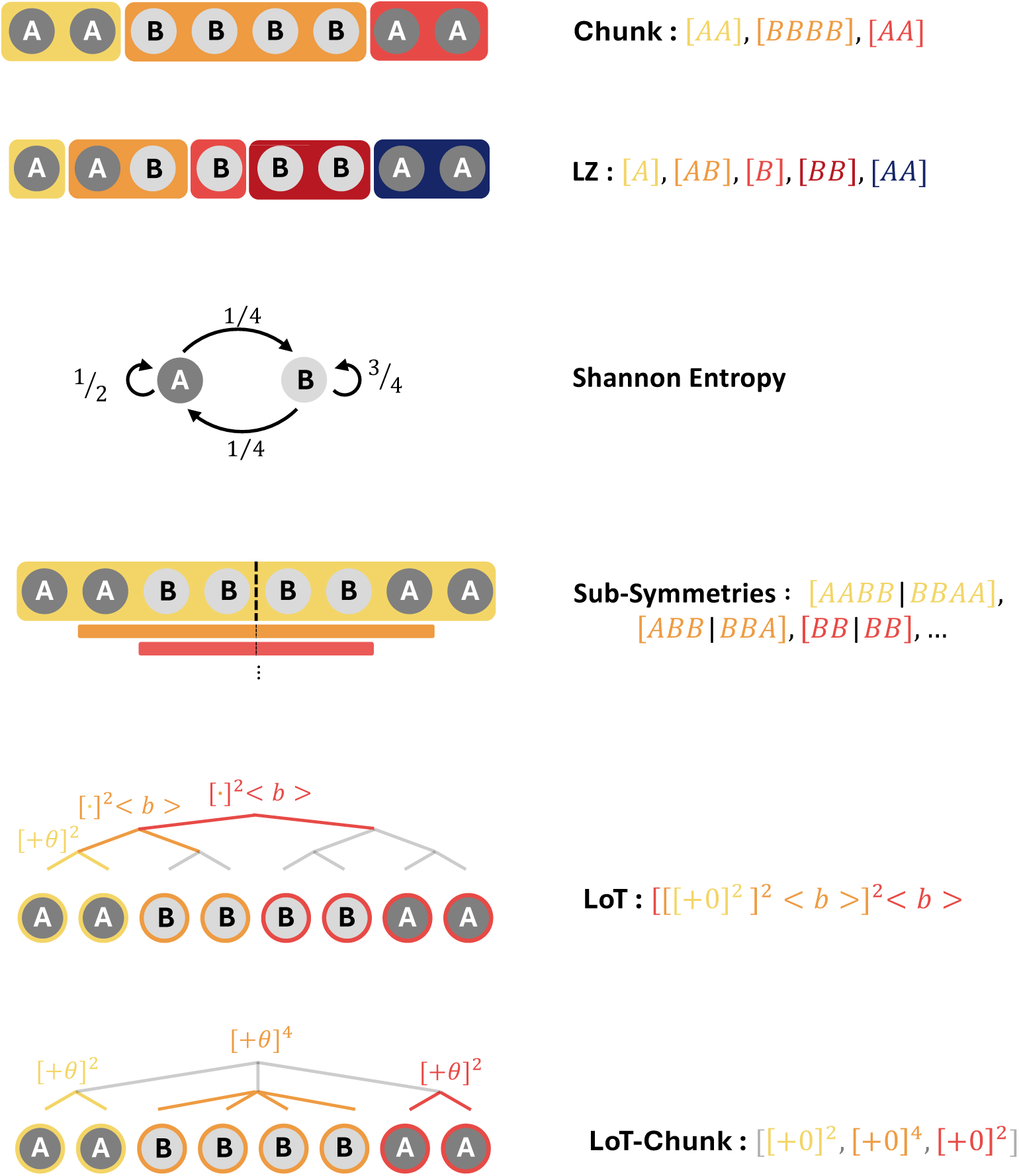
Illustrations of the competing models for pattern encoding.

